# Pathological α-synuclein elicits granulovacuolar degeneration independent of tau

**DOI:** 10.1101/2024.12.27.630547

**Authors:** Dylan J. Dues, Madalynn L. Erb, Alysa Kasen, Naman Vatsa, Erin T. Williams, An Phu Tran Nguyen, Michael X. Henderson, Darren J. Moore

## Abstract

**Background:** Pathologic heterogeneity is a hallmark of Lewy body dementia (LBD), yet the impact of Lewy pathology on co-pathologies is poorly understood. Lewy pathology, containing α-synuclein, is often associated with regional tau pathology burden in LBD. Similarly, granulovacuolar degeneration bodies (GVBs) have been associated with tau pathology in Alzheimer’s disease. Interestingly, GVBs have been detected in a broad range of neurodegenerative conditions including both α-synucleinopathies and tauopathies. Despite the frequent co-occurrence, little is known about the relationship between α-synuclein, tau, and granulovacuolar degeneration.

**Methods:** We developed a mouse model of limbic-predominant α-synucleinopathy by stereotactic injection of α-synuclein pre-formed fibrils (PFFs) into the basal forebrain. This model was used to investigate the relationship of α-synuclein pathology with tau and GVB formation.

**Results:** Our model displayed widespread α-synuclein pathology with a limbic predominant distribution. Aberrantly phosphorylated tau accumulated in a subset of α-synuclein inclusion-bearing neurons, often colocalized with lysosomes. Many of these same neurons also contained CHMP2b- and CK1δ-positive granules, established markers of GVBs, which suggests a link between tau accumulation and GVB formation. Despite this observation, GVBs were also detected in tau-deficient mice following PFF-injection, suggesting that pathological α-synuclein alone is sufficient to elicit GVB formation.

**Conclusions:** Our findings support that α-synuclein pathology can independently elicit granulovacuolar degeneration. The frequent co-accumulation of tau and GVBs suggests a parallel mechanism of cellular dysfunction. The ability of α-synuclein pathology to drive GVB formation in the absence of tau highlights the broader relevance of this process to neurodegeneration with relevance to the pathobiology of LBD.

## Background

In the spectrum of Lewy body dementia (LBD), the coexistence of Lewy pathology with Alzheimer’s disease neuropathologic change (ADNC) is the rule rather than the exception [1–5]. Although fewer Parkinson’s disease (PD) cases exhibit moderate to severe ADNC, this is drastically increased in cases of Parkinson’s disease dementia (PDD) and dementia with Lewy bodies (DLB) [6,7]. This trend suggests that the co-occurrence of Lewy pathology with AD-like tau pathology may be especially pertinent to the etiology of cognitive dysfunction in LBD [8]. Similarly, the presence of Lewy pathology in Alzheimer’s disease (AD) is common and is suspected to modify both motor and non-motor clinical features [9,10]. Thus, pathologic heterogeneity is a common feature of the α-synucleinopathies and may be highly relevant to the manifestation of both amnestic and non-amnestic cognitive features.

A major component of Lewy pathology is the small, pre-synaptic protein α-synuclein [11,12]. Both familial *SNCA* mutations (A53T, A30P, E46K) and *SNCA* gene multiplications are causative for LBD [13–16]. Intriguingly, the presence of comorbid tau pathology has been detected in these rare familial cases as well as in transgenic mouse lines expressing mutant forms of α-synuclein [17–22]. Post-mortem analysis of the LBD hippocampus has revealed that α-synuclein and tau pathologies follow overlapping, yet distinct, disease-associated subregional patterns [23,24]. The accumulation of multiple proteinopathies in dementia is associated with more severe cognitive impairment and accelerated decline, supporting an additive or possibly even synergistic interaction [1,7,25,26]. Despite this evidence, it is not clear how distinct neuropathologies arise and coalesce in the context of aging and LBD. It is also plausible that tau contributes to the progression of Lewy pathology and LBD-related cognitive dysfunction independent of neurofibrillary tangle formation. Tau epitopes have been detected in Lewy bodies by double immunofluorescence and immunohistochemistry, providing evidence that the formation of Lewy pathology incites cellular changes that support the aberrant accumulation of phosphorylated tau [27,28].

Here, we present a novel model of forebrain α-synucleinopathy in mice to study to induction of additional cellular pathologies. Following stereotactic injection of mouse α-synuclein pre-formed fibrils (PFFs) into the basal forebrain, wild-type mice exhibit widespread and bilateral formation of templated α-synuclein pathology. Inclusion burden was assessed at 3 months post-injection and revealed a limbic-predominant distribution. Both neuritic and cell body inclusions were readily detected by immunostaining for pS129-α-synuclein. Beyond the formation of α-synuclein pathology, we identified the parallel accrual of phosphorylated tau clustered around α-synuclein inclusions. Although, many regions with α-synuclein inclusions showed a similar pattern of hyperphosphorylated tau accumulation, there were clear exceptions that had abundant α-synuclein pathology but no hyperphosphorylated tau. Granule neurons of the dentate gyrus harbored few, if any, tau puncta, while pyramidal neurons in the hippocampal CA1 subfield have extensive granular tau deposits. Rather than a tangle-like structure, these puncta were morphologically reminiscent of tau accumulation previously observed alongside granulovacuolar degeneration bodies (GVBs). GVBs are poorly understood organelles that are often observed in the hippocampal CA1 subfield in AD [29,30]. GVBs have traditionally been found to coalesce with pre-tangle tau assemblies, rather than neurofibrillary tangles. Experimentally, GVBs have been demonstrated to form in the mouse hippocampus following application of synthetic human K18 tau P301L pre-formed fibrils [31]. More recently, GVBs have been shown to accumulate in primary neurons following application of α-synuclein PFFs, suggesting that GVB formation may be triggered by either tau or α-synuclein aggregation [32]. Post-mortem assessments support that multiple neurodegenerative disease classifications harbor granulovacuolar degeneration, and that this process may represent a fundamental neuronal response to protein aggregation and endolysosomal dysfunction [33]. However, it remains uncertain whether the formation of GVBs is predominantly a protective or degenerative process [34].

Here, for the first time *in vivo*, we identify the formation and limbic enrichment of GVBs as a response to templated α-synuclein pathology in the mouse forebrain. These intraneuronal granular assemblies co-label with CHMP2b and CK1δ, established markers of granulovacuolar degeneration [35]. CHMP2b- and CK1δ-positive granules were scattered within CA1 pyramidal neurons, adorning tendril-like α-synuclein inclusions and overlapping with phospho-tau puncta. These findings suggest a relationship between GVB formation and tau accumulation following α-synuclein aggregation, though we also demonstrate that GVBs accrue in the absence of endogenous tau. While granulovacuolar degeneration has historically been associated with tauopathy, our data indicate that it may also be triggered by forebrain α-synucleinopathy. Given observations of concomitant α-synuclein and tau pathology in the context of LBD-related cognitive dysfunction, the impact of GVB formation on neurodegeneration remains to be clarified. Further studies into the molecular basis of this convergence are thus warranted.

## Results

### Basal forebrain injection with α-synuclein PFFs elicits widespread α-synuclein pathology

We have previously examined intra-hippocampal injection of α-synuclein PFFs on the formation of α-synuclein pathology in the mouse hippocampus [36]. Under this paradigm, we observed that the CA2/3 subfield was especially susceptible to α-synuclein pathology and subsequent neurodegeneration, an observation that is consistent in the PFF model [37]. The susceptibility of the hippocampal CA2/3 subfield has also been demonstrated in studies of Lewy pathology in post-mortem LBD cases [38,39]. However, an important caveat of this model is that stereotactic injection directly into the hippocampus may influence pathologic change via mechanical injury and PFF exposure to resident non-neuronal cell types. Thus, we sought to develop a complementary mouse model of hippocampal α-synucleinopathy based on connectivity within the mouse brain without the need to directly target the hippocampus. We pursued stereotactic injection into the basal forebrain, specifically in the medial septal area and vertical limb of the diagonal band (**Figure 1A**). These regions exhibit strong connectivity across the limbic connectome and, critically, are distal to the hippocampal formation. Thus, PFF injection at this site allows for the induction of hippocampal “Lewy-like” pathology without the caveats of intra-hippocampal injection. We performed stereotactic injections in adult C57BL/6J mice followed by histopathologic analyses at 3 months post-injection (MPI). We sought to examine cellular changes prior to the onset of neurodegeneration. Under this paradigm, α-synuclein inclusion burden is induced with minimal neurodegenerative changes being detected at 3 MPI (**Figure S1**). Minor silver-positive neuritic degeneration was detected in the hippocampal fimbria and amygdala, but in the absence of any detectable cognitive or anxiety-related deficits, as measured by elevated plus maze, Y-maze, Barnes maze, and contextual fear conditioning (**Figure S1 A-B, Figure S2 A-I**).

**Figure 1.**
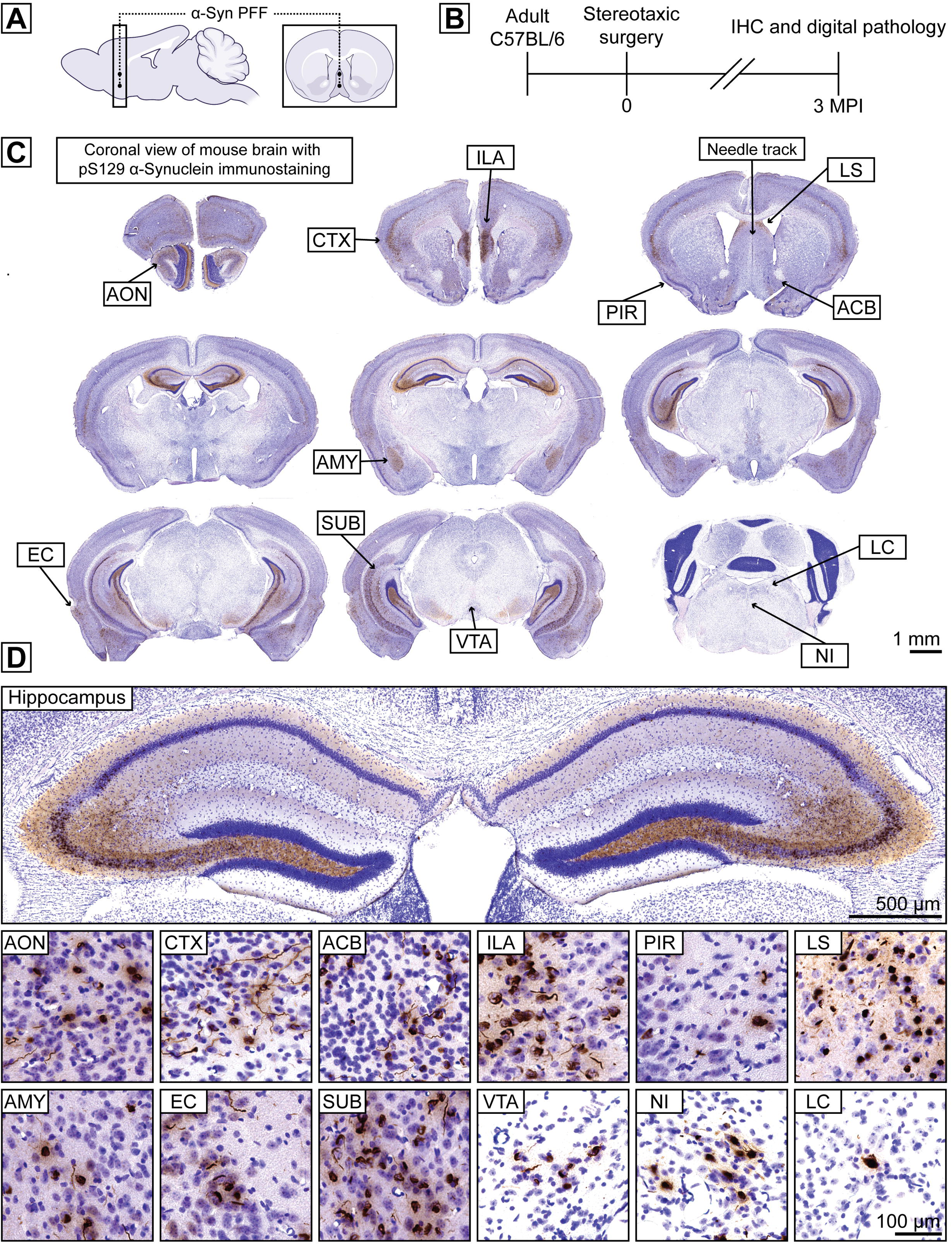
Basal forebrain injection with α-synuclein PFFs elicits limbic-predominant α-synuclein pathology. **A)** Representative schematic including sagittal and coronal view of α-synuclein PFF injection in a midline 2-point injection to the mouse basal forebrain. **B)** Timeline of stereotactic surgery in 2- to 3-month-old C57BL/6J mice followed by immunohistochemistry and digital pathology analyses at 3 months post-injection. **C)** Representative mouse coronal brain sections displaying pS129-α-synuclein immunostaining with Nissl counterstain. The needle track may be observed in the coronal section placed in the upper right quadrant. Scale bar: 1 mm. **D)** Higher magnification images of various regions from mouse coronal brain sections displayed in (**C**), the hippocampus exhibits bilateral α-synuclein pathology and inclusions are readily observed across the rostral-caudal axis. Abbreviations are as follows: anterior olfactory nucleus (AON), frontal cortex (CTX), nucleus accumbens (ACB), infralimbic area (ILA), piriform cortex (PIR), lateral septum (LS), amygdala (AMY), entorhinal cortex (EC), subiculum (SUB), ventral tegmental area (VTA), nucleus incertus (NI), locus coeruleus (LC). Scale bar: 500 μm for hippocampal image, 100 μm all other images.

We observed extensive α-synuclein pathology at 3 MPI, revealed by immunostaining for pS129-α-synuclein, a well-established and selective marker of aggregated α-synuclein (**Figure 1C**). The bulk of α-synuclein pathology was confined to limbic structures, signifying that basal forebrain injection induces limbic-predominant α-synucleinopathy. We observed bilateral pathology with good symmetry, which was particularly evident in the dorsal hippocampus and basolateral amygdala (**Figure 1D**). Pathological inclusions were detected in both neuronal cell bodies and processes and displayed a morphology comparable to what has been observed previously [40]. While many regions harbored α-synuclein pathology, this model revealed an amplified inclusion burden in a few select areas (**Figure 1D**). There was notably less pathology observed at the site of injection relative to more distal regions (**Figure S3 A-C**). The amygdala harbored an extensive inclusion burden, comparable to what has been observed under the intra-striatal paradigm. Abundant pathology was also present in the medial prefrontal cortex and adjacent forebrain cortical regions. A small number of inclusions were observed in select hindbrain regions such as the locus coeruleus, supporting a capacity for seeding across considerable distances (**Figure 1D**). Most apparent, however, was the distribution of α-synuclein pathology across the hippocampal axis (**Figure S4 A-E**). The pattern was similar to pathology patterns observed following intra-hippocampal injection [36]. However, we found that α-synuclein pathology in the basal forebrain model was much more broadly distributed, with greater involvement of the full rostral-caudal axis of the hippocampus. There was also less prominent involvement of the dentate gyrus, which is more in line with pathology in LBD brains [23]. Overall, these results support the use of basal forebrain injection to model limbic-predominant α-synuclein pathology in the mouse brain. It also avoids the confounding influence of extracellular α-synuclein PFFs which may act as an inflammatory modulator within the hippocampus.

### Accumulation of microtubule-associated protein tau occurs secondary to α-synuclein pathology and is enriched in the hippocampal CA1 subfield

This mouse model develops a unique limbic-predominant distribution of α-synuclein pathology, reminiscent of subfield patterns of Lewy pathology observed in the LBD hippocampus. Given our interest in whether Lewy pathology might influence other disease-relevant proteins associated with cognitive dysfunction, we sought to examine the impact of α-synuclein pathology on the microtubule-associated protein tau. We examined the posterior (ventral) portion of the hippocampus where α-synuclein inclusions were observed in both the dentate gyrus and CA1 subfield (**Figure 2**). Notably, across matched coronal brain sections we detected granular-appearing phosphorylated tau (AT8) puncta localizing to the soma of CA1 pyramidal neurons, but not in granule neurons of the dentate gyrus (**Figure 2**). This suggests that the cytosolic accumulation and aberrant phosphorylation of tau secondary to α-synuclein pathology may be influenced by neuronal phenotype. To better understand the distribution of α-synuclein and tau within the hippocampus, we quantified the subregional burden of α-synuclein pathology and accumulated tau puncta across the anterior-posterior hippocampal axis. The dentate gyrus exhibited substantially fewer α-synuclein inclusions with a slight increase in the posterior portion (**Figure 3A-B**). The CA2/3 subfield revealed an opposing pattern, in which α-synuclein pathology was anteriorly prominent (in the dorsal hippocampus) and reduced posteriorly (**Figure 3A-B**). Examination of the CA1/subiculum pyramidal layer revealed fewer inclusions in portions of the dorsal hippocampus, but drastically increased inclusions towards the intermediate-posterior hippocampus. Quantification revealed a similar pattern of phosphorylated tau puncta across the hippocampal axis, but with a more prominent burden in posterior sections of the CA1 subfield (**Figure 3C-D**). Qualitatively, α-synuclein pathology tended to concentrate in the distal segment of the intermediate hippocampal portion of the CA1 subfield along with greater enrichment in the superficial pyramidal layer (**Figure S4**). This pattern was also reflected in tau immunostained sections, with the transition from intermediate to ventral CA1 accounting for the most substantially affected portion of the hippocampus. Thus, our results depict the distribution and burden of phosphorylated tau as closely reflecting that of α-synuclein. Moreover, we identify the intermediate to posterior portions of the CA1 subfield as a hotspot in this model for concomitant α-synuclein inclusions and phosphorylated tau puncta (**Figure S5**).

**Figure 2.**
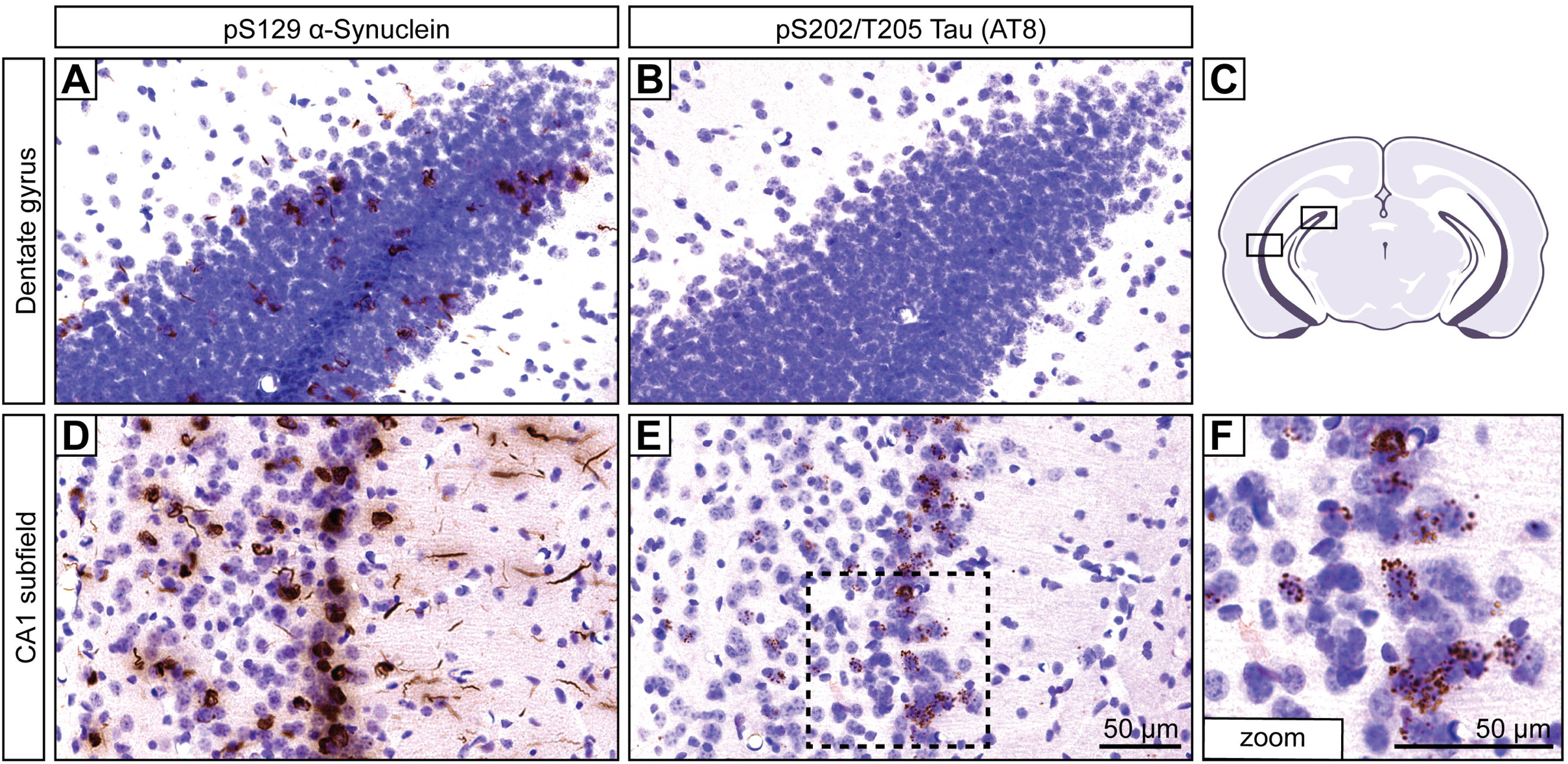
Aberrant accumulation of microtubule-associated protein tau is highly enriched in select hippocampal subfields of PFF-injected mice. **A-F)** Representative histological images displaying either pS129-α-synuclein (**A, D**) or pS202/T205-tau (AT8; **B, E-F**) immunostaining with Nissl counterstain. **A)** Sparse α-synuclein inclusions are observed in the dentate gyrus. **B)** Negligible tau puncta are observed in the dentate gyrus. **C)** Representative schematic showing the regions of interest in a mouse coronal section featuring the posterior hippocampus. **D)** Abundant α-synuclein inclusions are observed in the CA1 subfield. **E)** Tau puncta are readily observed in the CA1 subfield. Scale bar: 50 μm. **F)** Higher magnification inset from **(E)** reveals abundant tau puncta in a perisomatic distribution displaying a non-tangle-like morphology. Scale bar: 50 μm.

**Figure 3.**
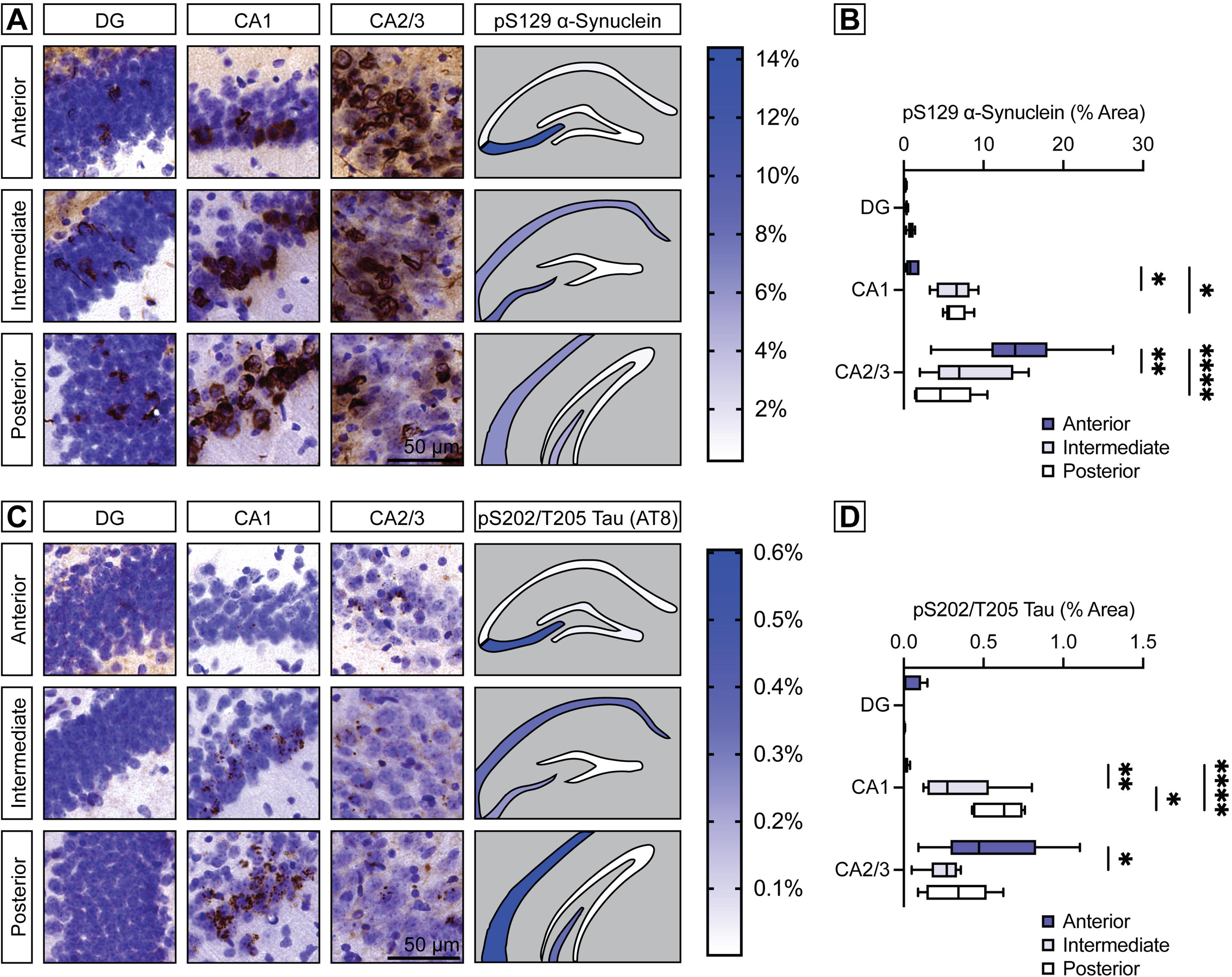
α-Synuclein and tau follow a similar subregional distribution across the hippocampal axis. **A)** Representative histological images displaying pS129-α-synuclein immunostaining with Nissl counterstain across the anterior-posterior axis of the hippocampus including sections of the dentate gyrus and CA1-3 subfields. A hippocampal heatmap displays the mean % area occupied of pS129-α-synuclein immunostaining by region. Scale bar: 50 μm. **B)** Quantification of % area occupied of pS129-α-synuclein immunostaining across the anterior-posterior axis of the hippocampus including sections of the dentate gyrus and CA1-3 subfields (box and whiskers plot with boxes defining the 25^th^ to 75^th^ percentiles and whiskers displaying the min-max value, *n* = 6 mice). \**P*<0.05, \*\**P*<0.01, or \*\*\*\**P*<0.0001 by two-way ANOVA with Tukey’s multiple comparisons test, as indicated. **C)** Representative histological images displaying pS202/T205-tau (AT8) immunostaining with Nissl counterstain across the anterior-posterior axis of the hippocampus including sections of the dentate gyrus and CA1-3 subfields (*n* = 6 mice). A hippocampal heatmap displays the mean % area occupied of pS202/T205-tau (AT8) immunostaining by region. Scale bar: 50 μm. **D)** Quantification of % area occupied of pS202/T205-tau (AT8) immunostaining across the anterior-posterior axis of the hippocampus including sections of the dentate gyrus and CA1-3 subfields (box and whiskers plot, *n* = 6 mice). \**P*<0.05, \*\**P*<0.01, or \*\*\*\**P*<0.0001 by two-way ANOVA with Tukey’s multiple comparisons test, as indicated.

### Pathological α-synuclein inclusions are adorned by granulovacuolar degeneration bodies and mislocalized tau

Given the interesting regional patterning of α-synuclein and tau across the hippocampal axis, we next sought to identify the intracellular localization of the two proteins via co-immunofluorescence. We focused our examination on the CA1 subfield of the intermediate to posterior portion of hippocampus, where α-synuclein and tau accumulation was enriched (**Figure S5**). Intriguingly, we found that pS129-α-synuclein and pS202/T205-tau (AT8) immunostaining identified overlapping but discrete structures within the soma of CA1 pyramidal neurons (**Figure 4**). Most granular tau puncta cluster within tendril-like α-synuclein inclusions resembling a nested configuration. In addition, the number of tau puncta per neuron varied considerably. This pattern suggests that tau accumulation is not an obligatory consequence of α-synuclein aggregation, but rather a reaction influenced by the underlaying variation of the neuronal response to α-synuclein. Notably, assessment of tau at an earlier time-point (1.5 MPI) suggests that tau puncta are not an immediate response to α-synuclein accumulation and that this process may be dynamic (**Figure S6**).

**Figure 4.**
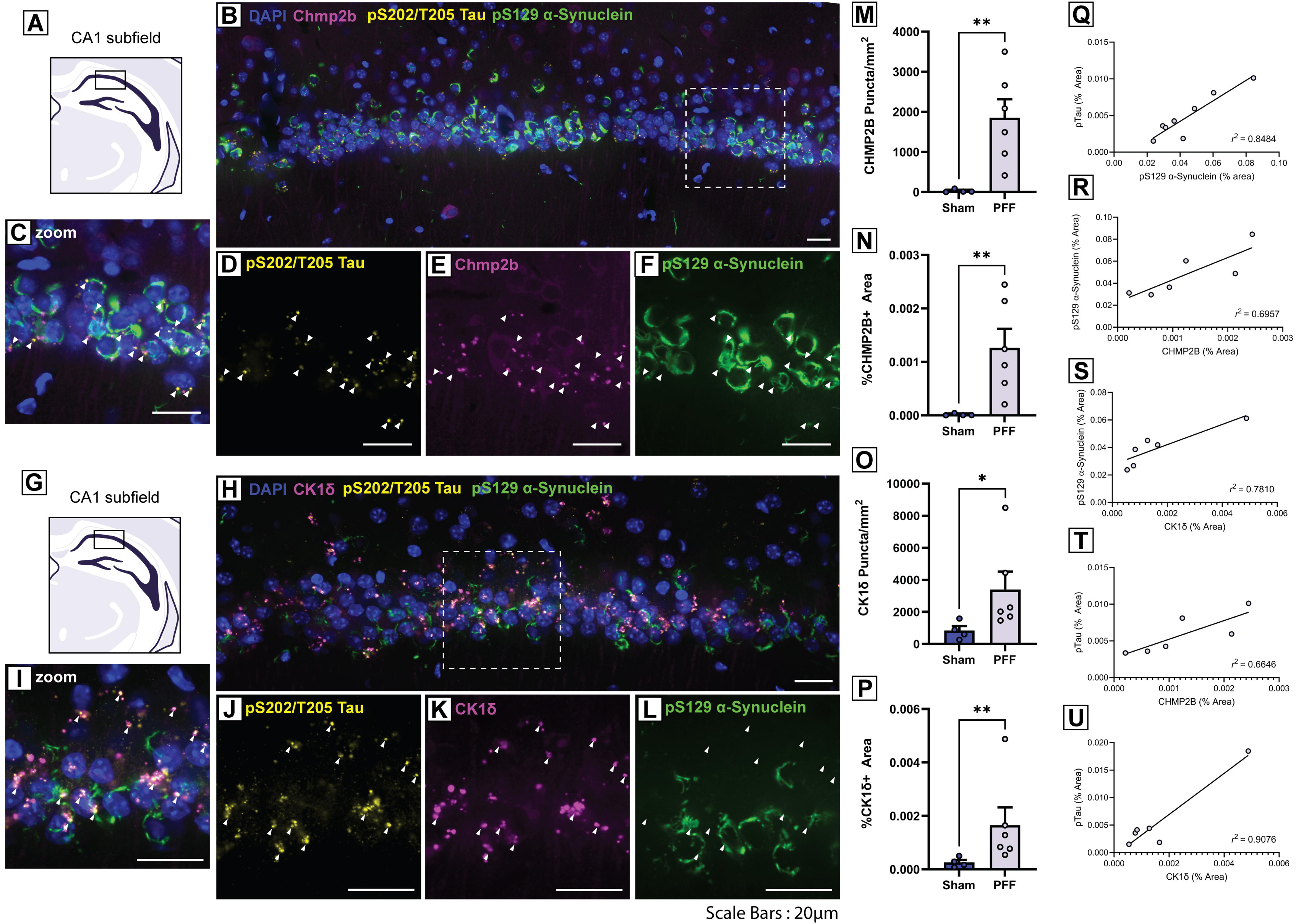
Cellular localization of α-synuclein, tau, and granulovacuolar degeneration bodies in the distal CA1 subfield. **A)** Representative schematic of the distal CA1 pyramidal layer of the intermediate/posterior hippocampus. **B)** Confocal immunofluorescent image showing pS129-α-synuclein, pS202/T205-tau, and CHMP2b in the CA1. **C-F)** Inset image from **(B)** showing **(C)** combined, **(D)** pS202/T205-tau, **(E)** CHMP2b, and **(F)** pS129-α-synuclein. Arrows indicate pS202/T205-tau inclusions. **G)** Representative schematic of the distal CA1 pyramidal layer of the intermediate/posterior hippocampus. **H)** Confocal immunofluorescent image showing pS129-α-synuclein, pS202/T205-tau, and CK1δ in the CA1. **I-L)** Inset image from **(H)** showing **(I)** combined, **(J)** pS202/T205-tau, **(K)** CK1δ, and **(L)** pS129-α-synuclein. Arrows indicate pS202/T205-tau inclusions. **M-P)** Density of **(M-N)** CHMP2b or **(O-P)** CK1δ granules in the CA1 (mean ± SEM, *n* = 4-6 images from 2-3 mice/group). Graphs show the number of **(M)** CHMP2b or **(O)** CK1δ granules/mm^2^ or the % area occupied by **(N)** CHMP2b or **(P)** CK1δ (*n* = 4-6 hippocampal sections). \**P*<0.05, or \*\**P*<0.01 by unpaired Student’s *t*-test. **Q-U)** Graphs show the relationship of % area occupied by pS202/T205-tau, pS129-α-synuclein, CHMP2b or CK1δ inclusions in CA1. Scale bars: 20 μm.

Since pre-tangle tau pathology in the hippocampal CA1 subfield has been associated with the presence of granulovacuolar degeneration bodies in AD, we concurrently probed for the established GVB markers CHMP2b and CK1δ. We identified accumulation of large CHMP2b-positive granules in α-synuclein inclusion-bearing neurons (**Figure 4A-F**). We also detected the formation of CK1δ-positive granules in inclusion-bearing neurons (**Figure 4G-L**). Importantly, the accumulation of CHMP2b- and CK1δ-positive granules in fibril-injected mice were wholly distinct from the more diffuse staining observed in the soma of PBS-injected mice (**Figure S7**). Our results point towards a pathologic cascade whereby the formation of α-synuclein pathology elicits the concurrent accrual of tau and granulovacuolar degeneration. Of note, intra-hippocampal PFF-injected mice revealed greater involvement of the dentate gyrus, with consistently minimal co-accrual of tau puncta and GVBs despite substantial α-synuclein burden (**Figure S8 A-F**). This suggests that these secondary features may be convergent, or at least similarly influenced by neuronal phenotype as cortical regions in the hippocampal paradigm also yielded α-synuclein, tau, and GVB formation (**Figure S8 G-L**).

### Endogenous tau is dispensable for the formation of GVBs induced by α-synuclein pathology

Given the long-standing association between tauopathy and granulovacuolar degeneration, we sought to determine whether the presence of endogenous tau was a requirement for the formation of GVBs *in vivo* following application of α-synuclein PFFs. We first observed that endogenous tau is not required for the templated formation of α-synuclein pathology in mice, in agreement with previous studies [41]. In wild-type mice, most inclusion-bearing neurons that harbored GVBs also exhibited tau accumulation, suggesting that these processes may be inter-related. Remarkably, we identified CHMP2b-positive GVBs associated with α-synuclein inclusions in the hippocampus of *Mapt* knockout mice following intra-hippocampal injection with α-synuclein PFFs (**Figure 5**). This data supports that GVBs arise following the *in vivo* aggregation of α-synuclein independent of tau. It remains unclear whether the co-accumulation of tau in wild-type mice occurs independent of GVB formation, as tau and GVB markers both accrue in inclusion-bearing neurons (**Figure 5**). It may be the case that tau accumulation occurs in parallel to GVB formation, arising from endolysosomal pathway dysfunction. Alternatively, GVB-related changes may more directly impact tau, though additional studies are necessary to confirm whether an interaction between tau and GVB components occurs in this model.

**Figure 5.**
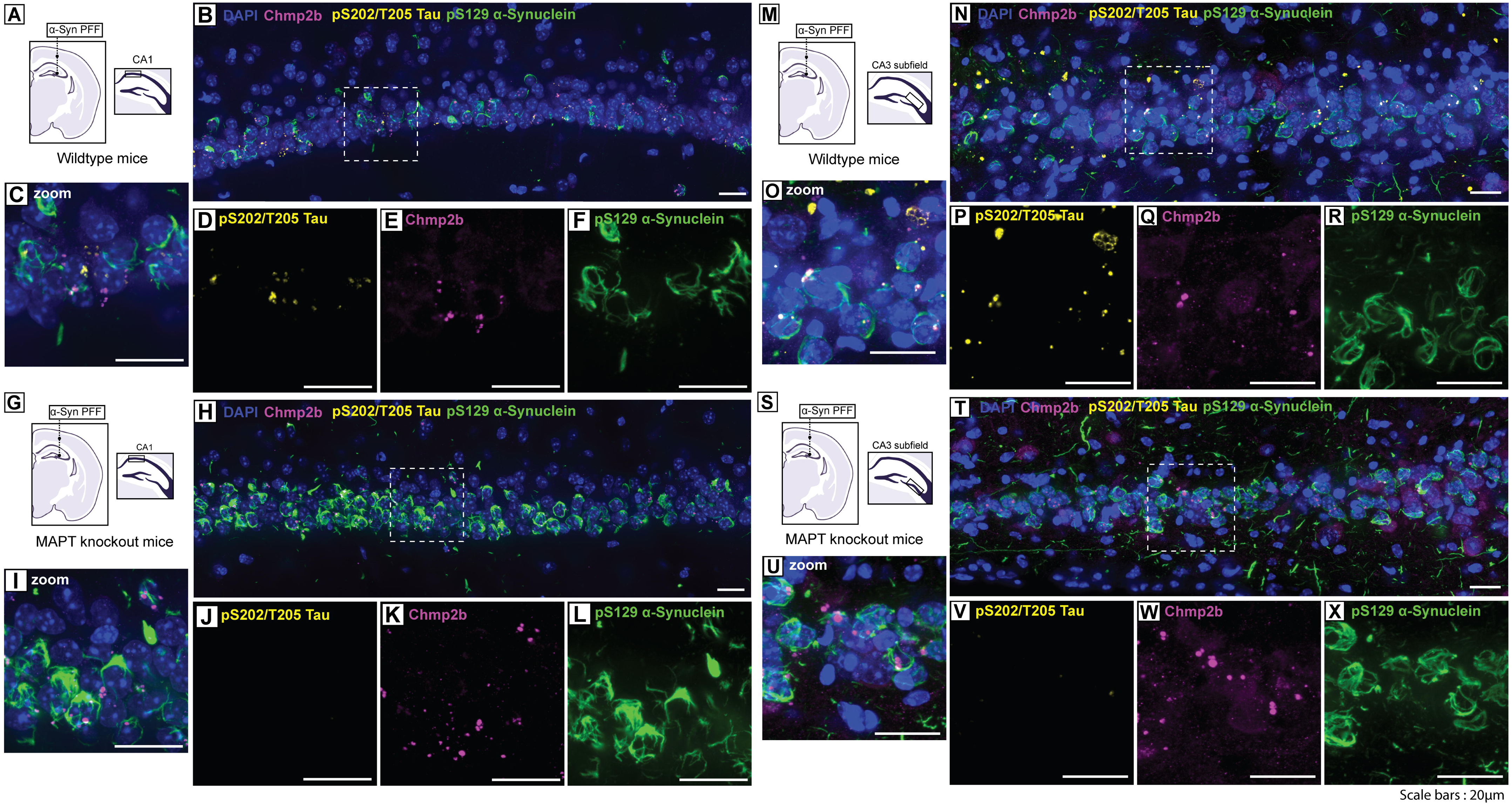
Endogenous microtubule-associated protein tau is not required for the formation of granulovacuolar degeneration bodies in the hippocampal PFF injection paradigm. **A)** Representative schematic of the hippocampal injection paradigm in wild-type mice. **B)** Confocal immunofluorescent image showing pS129-α-synuclein, pS202/T205-tau, and CHMP2b in the CA1 of wild-type mice. **C-F)** Inset image from **(B)** showing **(C)** combined, **(D)** pS202/T205-tau, **(E)** CHMP2b, and **(F)** pS129-α-synuclein. **G)** Representative schematic of the hippocampal injection paradigm in *MAPT* knockout mice. **H)** Confocal immunofluorescent image showing pS129-α-synuclein, pS202/T205-tau, and CHMP2b in the CA1 of *MAPT* knockout mice. **I-L)** Inset image from **(H)** showing **(I)** combined, **(J)** pS202/T205-tau, **(K)** CHMP2b, and **(L)** pS129-α-synuclein. **M)** Representative schematic of the hippocampal injection paradigm in wild-type mice. **N)** Confocal immunofluorescent image showing pS129-α-synuclein, pS202/T205-tau, and CHMP2b in the CA3 of wild-type mice. **O-R)** Inset image from **(N)** showing **(O)** combined, **(P)** pS202/T205-tau, **(Q)** CHMP2b, and **(R)** pS129-α-synuclein. **S)** Representative schematic of the hippocampal injection paradigm in *MAPT* knockout mice. **T)** Confocal immunofluorescent image showing pS129-α-synuclein, pS202/T205-tau, and CHMP2b in the CA3 of *MAPT* knockout mice. **U-X)** Inset image from **(T)** showing **(U)** combined, **(V)** pS202/T205-tau, **(W)** CHMP2b, and **(X)** pS129-α-synuclein. Scale bars: 20 μm.

### The confluence of α-synuclein and tau with granulovacuolar degeneration bodies is detected in primary hippocampal neurons treated with α-synuclein PFFs

After observing the induction of GVBs in the hippocampus *in vivo*, we next sought to evaluate GVBs in dissociated hippocampal neuron cultures, where effects of dose and individual cells could be characterized. Primary hippocampal neurons were exposed to α-synuclein PFFs at 7 days *in vitro* (DIV) and assessed at 21 DIV (**Figure 6**). Reflective of our observations *in vivo*, we identified the accumulation of cytosolic tau puncta and GVBs in neurons. Granular CK1δ-positive puncta decorated α-synuclein inclusions in neurons (**Figure 6A-C**). We next assessed the formation of CK1δ-positive GVBs following the application of varying concentrations of α-synuclein PFFs. Co-occurrence of GVBs with α-synuclein inclusions was concentration-dependent (**Figure 6D**). A similar dose dependence was also apparent in induction of phosphorylated tau puncta and CK1δ (**Figure 6E-H**). These findings point towards a cell-autonomous process by which α-synuclein pathology elicits GVB development.

**Figure 6.**
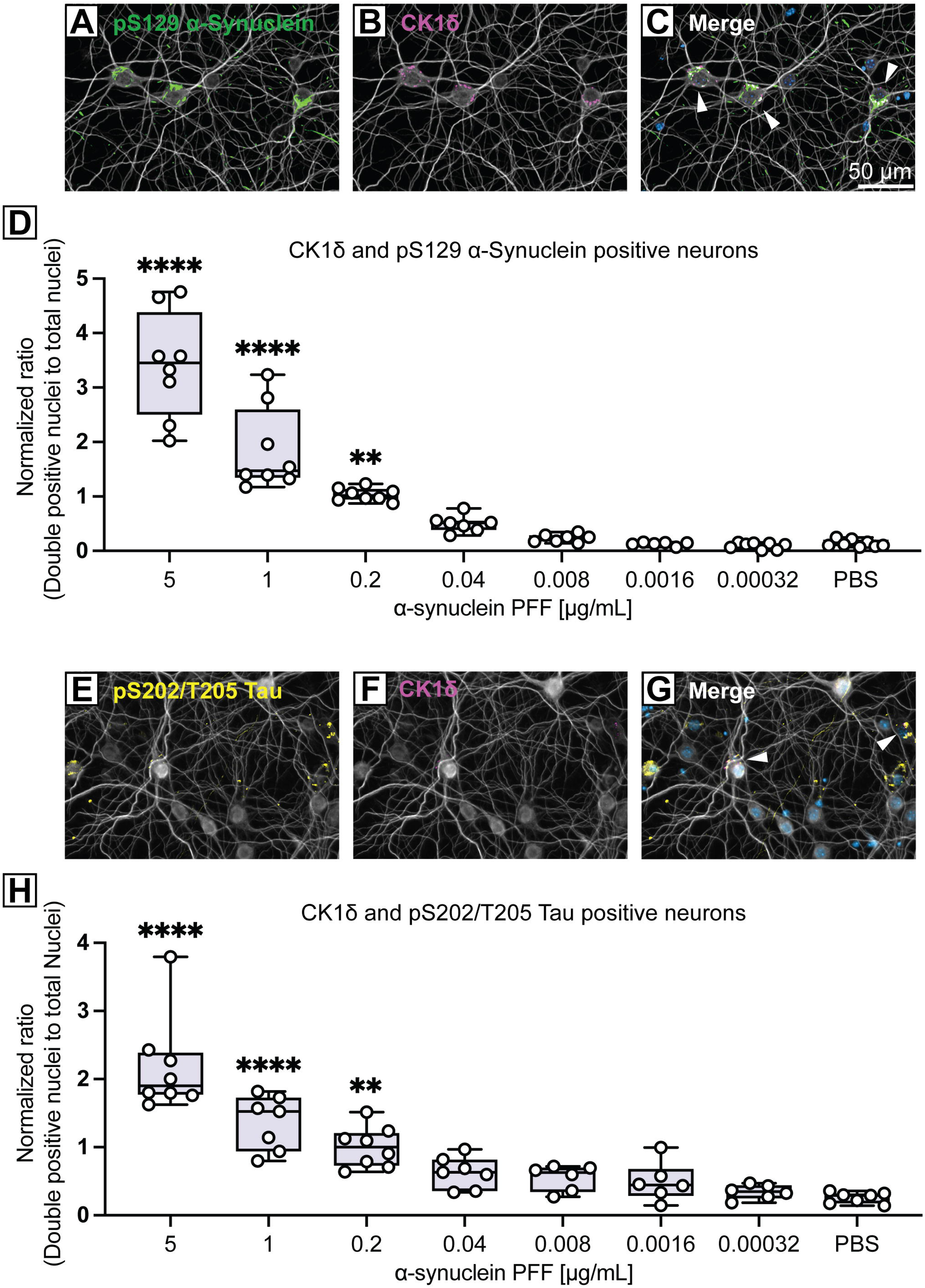
Co-accumulation of granulovacuolar degeneration bodies with α-synuclein inclusions in mouse primary hippocampal neurons is enhanced by α-synuclein PFF in a concentration-dependent manner. **A-C)** Immunofluorescent image of primary hippocampal neurons treated with α-synuclein PFFs stained for **(A)** pS129-α-synuclein, **(B)** CK1δ, and **(C)** combined with DAPI. Scale bar: 50 μm. **D)** Quantification of double-positive neurons for both pS129-α-synuclein and CK1δ across a range of α-synuclein PFF concentrations (µg/ml)) including PBS control (mean ± SEM, *n* = 9 (3 replicates per culture and 3 cultures/condition)). \*\**P*<0.01 or \*\*\*\**P*<0.0001 by one-way ANOVA with Dunnett’s test, as indicated. **E-G)** Immunofluorescent image of primary hippocampal neurons treated with α-synuclein PFFs stained for **(E)** pS202/T205-tau, **(F)** CK1δ, and **(G)** combined with DAPI. Scale bar: 50 μm. **H)** Quantification of double-positive neurons for both pS202/T205-tau and CK1δ across a range of α-synuclein PFF concentrations including PBS control (mean ± SEM, *n* = 9 (3 replicates per culture and 3 cultures/condition). \*\**P*<0.01 or \*\*\*\**P*<0.0001 by one-way ANOVA with Dunnett’s test, as indicated.

### Granular tau accumulates in lysosomes and is associated with changes in lysosomal morphology *in vivo*

To further explore the cellular response to pathological α-synuclein inclusions and tau granules we examined lysosomal morphology in affected brain regions including the CA1, CA3 and dentate gyrus of the hippocampus and the amygdala (**Figure 7 and S9**). In the hippocampus and amygdala tau granules colocalize with LAMP2-positive lysosomes (**Figure 7**). Additionally, lysosomes that are associated with tau granules are larger than lysosomes in neighboring cells without tau granules or lysosomes in sham mice (**Figure 7**). Lysosomal swelling is a signifier of lysosomal stress, which is consistent with previous reports showing that accumulation of pathological tau causes lysosomal permeabilization and deacidification [42]. Notably, we do not observe lysosomal swelling in the dentate gyrus (**Figure S9**), where pathological α-synuclein inclusions occur without tau granule formation. Transcription Factor E3 (TFE3) can respond to certain types of lysosomal or nutrient stress by upregulating lysosomal biogenesis genes. However, lysosomal swelling is not accompanied by nuclear translocation of TFE3 in PFF-injected mice (**Figure S10**).

**Figure 7.**
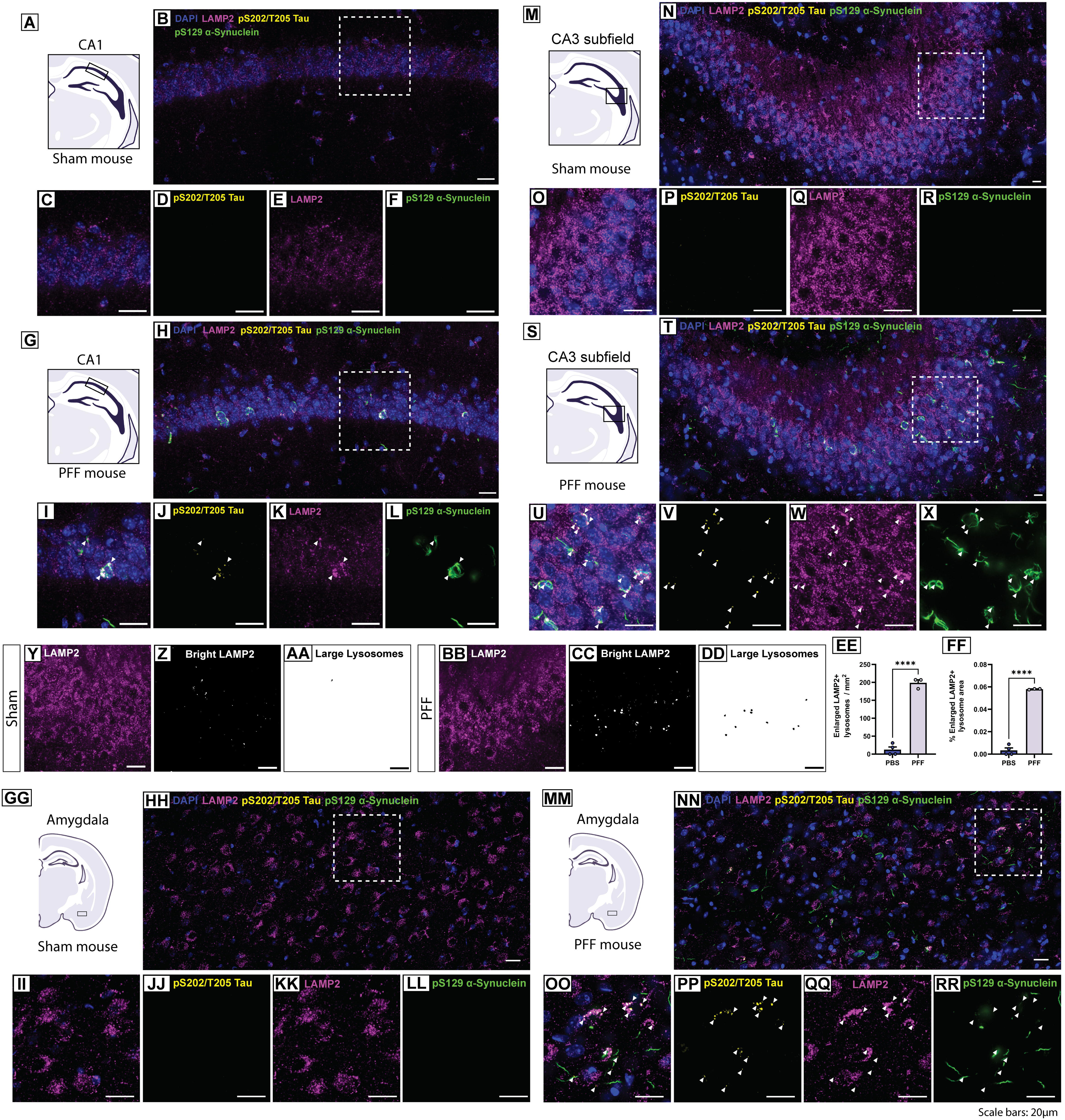
pS202/T205-tau inclusions are associated with enlarged lysosomes. **A, G)** Representative schematic of the distal CA1 pyramidal layer of the intermediate/posterior hippocampus in **(A)** Sham or **(G)** PFF-injected mice. **B)** Confocal immunofluorescent image of pS129-α-synuclein, pS202/T205-tau, and LAMP2 in the CA1 of Sham mice. **C-F)** Inset image from **(B)** showing **(C)** combined, **(D)** pS202/T205-tau, **(E)** LAMP2, and **(F)** pS129-α-synuclein. **H)** Image of pS129-α-synuclein, pS202/T205-tau, and LAMP2 in the CA1 of PFF-injected mice. **I-L)** Inset image from **(H)** showing **(I)** combined, **(J)** pS202/T205-tau, **(K)** LAMP2, and **(L)** pS129-α-synuclein. Arrows indicate pS202/T205-tau inclusions. **M, S)** Representative schematic of the CA3 of the intermediate/posterior hippocampus in **(M)** Sham or **(S)** PFF-injected mice. **N)** Image of pS129-α-synuclein, pS202/T205-tau, and LAMP2 in the CA3 of Sham mice. **O-R)** Inset image from **(N)** showing **(O)** combined, **(P)** pS202/T205-tau, **(Q)** LAMP2, and **(R)** pS129-α-synuclein. **T)** Image of pS129-α-synuclein, pS202/T205-tau, and LAMP2 in the CA3 of PFF mice. **U-X)** Inset image from **(T)** showing **(U)** combined, **(V)** pS202/T205-tau, **(W)** LAMP2, and **(X)** pS129-α-synuclein. Arrows indicate pS202/T205-tau inclusions. **Y-FF)** Quantification of enlarged lysosomes in the CA3. **Y, BB)** LAMP2 immunofluorescence in the CA3 of **(Y)** Sham or **(BB)** PFF mice. **Z, CC)** Brightest LAMP2 puncta in Sham or PFF mice. **AA, DD)** Large LAMP2-positive lysosomes in Sham or PFF mice. Graphs show **(EE)** the number of enlarged LAMP2+ lysosomes / mm^2^ or **(FF)** the % area of enlarged lysosomes (mean ± SEM, *n* = 3-4 mice/group). \*\*\*\**P*<0.0001 by unpaired Student’s *t*-test. **(GG, MM)** Representative schematic of the amygdala in **(GG)** Sham or **(MM)** PFF mice. **HH)** Confocal immunofluorescent image showing pS129-α-synuclein, pS202/T205-tau, and LAMP2 in the amygdala of Sham mice. **II-LL)** Inset image from **(HH)** showing **(II)** combined, **(JJ)** pS202/T205-tau, **(KK)** LAMP2, and **(LL)** pS129-α-synuclein. **NN)** Image of pS129-α-synuclein, pS202/T205-tau, and LAMP2 in the amygdala of PFF mice. **(OO)** Inset image from **(NN)** showing **(OO)** combined, **(PP)** pS202/T205-tau, **(QQ)** LAMP2, and **(RR)** pS129-α-synuclein. Arrows indicate pS202/T205-tau inclusions. Scale bars: 20 µm.

Enlarged lysosomes were detected in inclusion-bearing neurons that harbored tau granules (**Figure 7**). However, α-synuclein inclusions are most likely not engulfed by these organelles as their tendril-like structure is distinct from the vesicular morphology of lysosomes. Alternatively, we observe considerable overlap between pS129-α-synuclein and p62 in the CA1, the CA3 and the amygdala (**Figure 8**), which agrees with previous reports from experiments utilizing α-synuclein PFFs in cells or brain tissue [43–45]. Like pS129-α-synuclein, p62 inclusions were absent from Sham mice **(Figure S11)**. p62 immunostaining resembles the structure of α-synuclein inclusions and occurs with and without pS129-α-synuclein immunoreactivity in PFF mice, likely decorating α-synuclein inclusions with and without phosphorylated S129 (**Figure 8**). Notably, p62 structures frequently occur in neurons with tau granules but p62 does not share the granule morphology of pathological tau (**Figure 8**). p62 accumulation on α-synuclein inclusions indicates targeting of these proteinaceous aggregates for degradation through the autophagy-lysosomal pathway. Previous reports show that p62-labeled α-synuclein inclusions fail to be incorporated into autolysosomes and are subsequently not degraded by autophagy, which is consistent with the observation that pS129-α-synuclein inclusions are not engulfed by LAMP2-positive lysosomes (**Figure 7**) [43].

**Figure 8.**
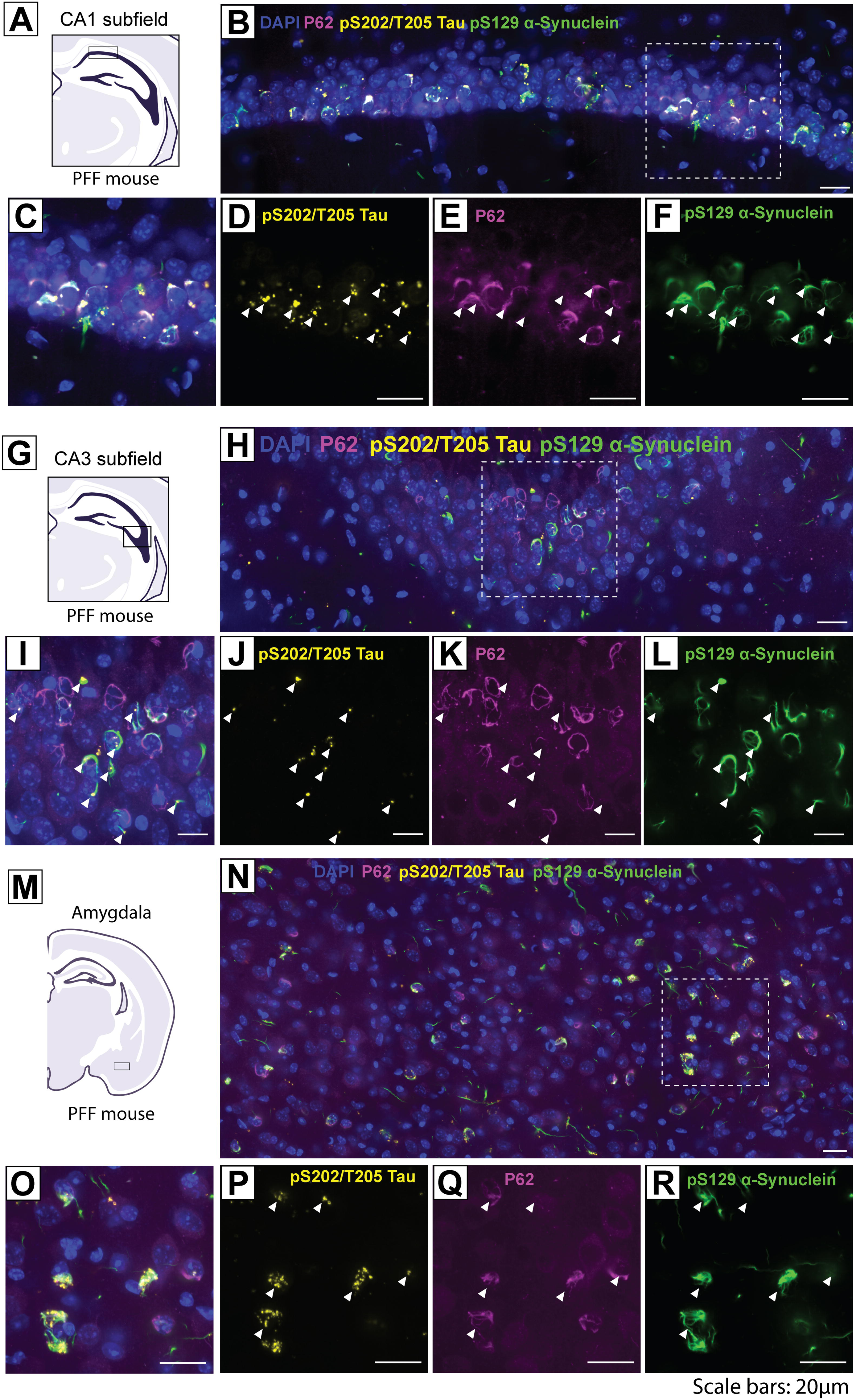
p62 inclusions are associated with pS129-α-synuclein pathology. **A, G, M)** Representative schematic of **(A)** the distal CA1 pyramidal layer of the intermediate/posterior hippocampus, **(G)** the CA3 of the intermediate/posterior hippocampus or **(M)** the amygdala in PFF-injected mice. **B, H, N)** Confocal immunofluorescent image showing pS129-α-synuclein, pS202/T205-tau, and p62 in the **B)** CA1, **H)** CA3 and **N)** amygdala. **C-F, I-L, O-R)** Inset image from **(B, H, N)** showing **(C, I, O)** combined, **(D, J, P)** pS202/T205-tau, **(E, K, Q)** p62, and **(F, L, R)** pS129-α-synuclein in **(C-F)** CA1, **(I-L)** CA3 and **(O-R)** amygdala. Arrows indicate pS202/T205-tau inclusions. Scale bars: 20 µm.

### Co-accumulation of GVBs with α-synuclein and tau in the hippocampus and amygdala of human LBD cases

We performed a histological analysis with cases of LBD, PDD or PD to probe the co-accumulation of GVBs, tau, and α-synuclein in the hippocampus and amygdala. We selected 3 cases with confirmed limbic Lewy pathology (**Figure S12**). In all cases, the presence of GVBs was initially identified through staining for CHMP2b and CK1δ (**Figure S13**). Notably, the GVB markers CHMP2b and CK1δ are present in neurons exhibiting accumulation of phosphorylated α-synuclein but are seemingly absent from neurons containing mature Lewy bodies (**Figure 9**). This observation parallels previous studies, where GVBs have been identified in neurons harboring pre-tangle tau accumulation rather than mature neurofibrillary tangles [35,46–48]. The appearance of GVBs in neurons with early-stage tau or α-synuclein accumulation suggests that their formation may represent a cellular stress response to initial misfolding and aggregation events, rather than a downstream consequence of mature inclusion formation.

**Figure 9.**
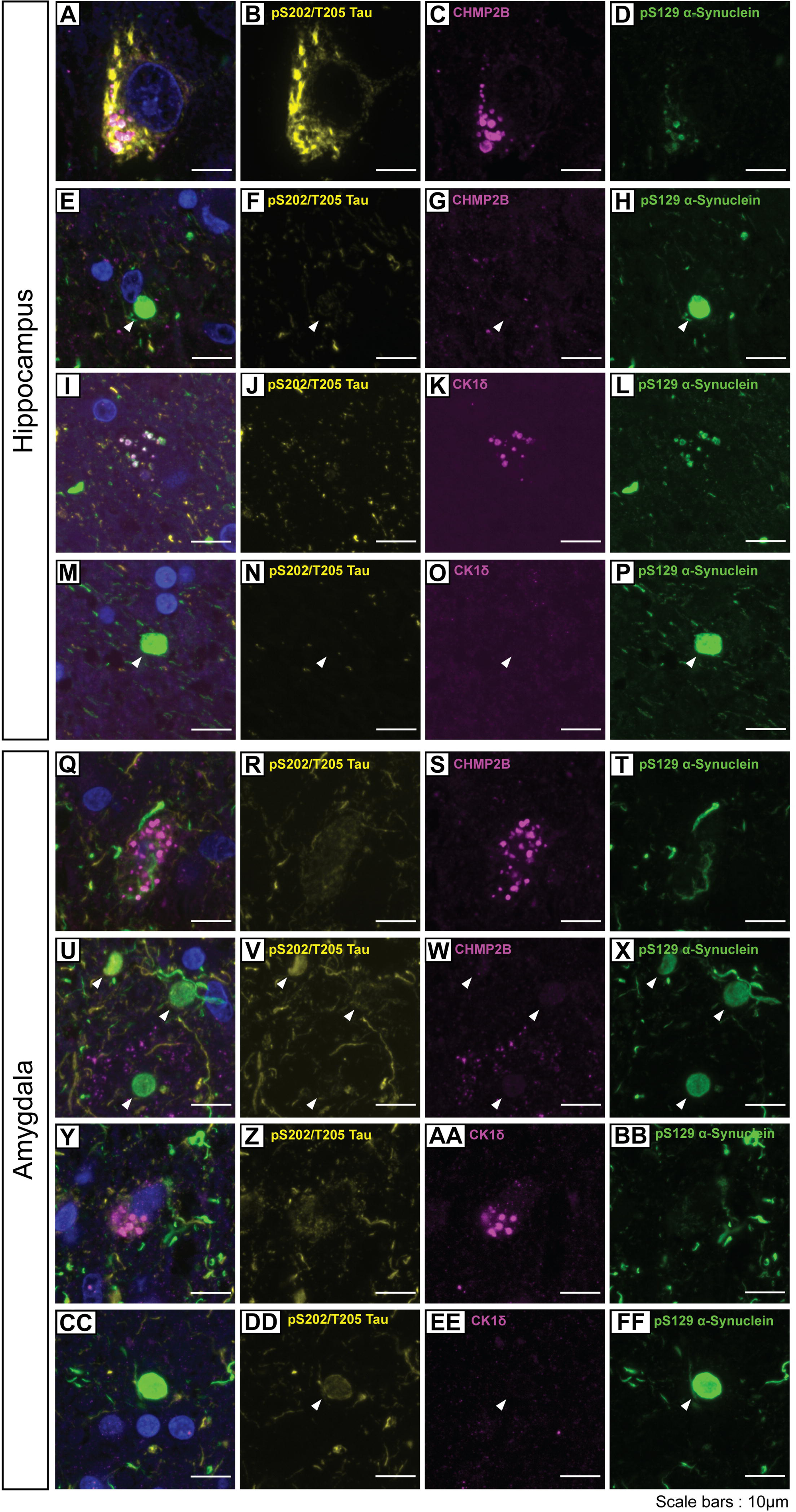
Human LBD brains depicting the confluence of α-synuclein, tau, and GVB markers. **A-P)** Representative confocal immunofluorescent images showing the staining patterns of pS129-α-synuclein, pS202/T205-tau, and CHMP2b **(A-H)** or CK1δ **(I-P)** in the CA1/CA3 subfields of the hippocampus. **Q-FF)** Representative confocal immunofluorescent images showing the staining patterns of pS129-α-synuclein, pS202/T205-tau, and CHMP2b **(Q-X)** or CK1δ **(Y-FF)** in the amygdala. Arrows indicate mature classical Lewy bodies that typically lack CHMP2b or CK1δ puncta. Representative images were derived from 3 LBD cases. Scale bars: 10 µm.

## Discussion

In this study we present a basal forebrain injection paradigm using α-synuclein PFFs to generate a novel model of limbic-predominant α-synuclein pathology. This model has specific technical advantages over the intra-hippocampal injection paradigm and generates a broader distribution of hippocampal α-synuclein pathology. Perhaps most beneficial, this alternative model elicits bilateral hippocampal pathology without exposing the local hippocampal environment to α-synuclein PFFs or mechanical injury. This provides a valuable complement to studies using intra-hippocampal injection, as exposure of non-neuronal populations in the hippocampus to fibrillar α-synuclein may impact aspects of disease-relevant biology, particularly regarding inflammation. In our study, we observed GVB formation and hyperphosphorylated tau accumulation under both injection paradigms, supporting that either paradigm may be applied in studying these changes.

The spread of α-synuclein pathology throughout the brain is postulated to follow defined patterns of connectivity influenced by neuronal phenotype [49]. Similarly, we hypothesize that neuronal phenotype might shape the subsequent response to pathologic α-synuclein. While the basis for this variation is undefined, we provide evidence that the abnormal accrual of phosphorylated tau secondary to α-synuclein pathology is differentially evoked across neuronal populations in the hippocampus. Alterations to tau were most abundant in pyramidal neurons located in the distal CA1 subfield, while granule neurons of the dentate gyrus were largely spared. We cannot exclude a threshold effect, by which a certain burden of α-synuclein pathology prompts subsequent accumulation of tau. This appears unlikely, however, given that many other brains regions with low to moderate α-synuclein burden still exhibit tau puncta formation, and we did not observe extensive tau accumulation under the intra-hippocampal paradigm where the dentate gyrus is more heavily affected relative to the basal forebrain paradigm.

There is a pressing need for the development of α-synucleinopathy models that recapitulate the pathophysiology of LBD-related cognitive dysfunction. Based on the stereotypic distribution of Lewy pathology in LBD, the involvement of forebrain structures appears to be a critical step in the neurodegenerative process. Further, the complex accrual of multiple pathologies, aside from α-synuclein inclusions, make pathologic heterogeneity an additional consideration. While the addition of multiple pathologies is linked to worse clinical status, the mechanistic basis of this phenomena is poorly understood, and it is uncertain whether mixed pathological changes arise independently or reciprocally. Our findings suggest that concomitant pathological changes may be interlinked, particularly for α-synuclein, tau, and granulovacuolar degeneration in the forebrain. Fibrillar α-synuclein has been shown to directly interact with tau and initiate tau aggregation via a cross-seeding event [50]. Congruently, strains of α-synuclein PFFs have been shown to initiate AD tangle-like tau pathology in mice [51]. More recently, the co-injection of α-synuclein fibrils with AD-extracts in mice has demonstrated that α-synuclein may influence the pathologic spread of tau [52]. Some PFF-based studies in tau knockout mice have shown that the absence of tau partly ameliorates neurodegeneration [53]. However, others have found that loss of endogenous tau does not substantially alter the formation of α-synuclein pathology *in vivo*, and that neurodegeneration is not impacted by tau genotype [41]. Some of these discrepancies may be explained by a limited α-synuclein and tau interaction that is dependent on neuronal phenotype, as these studies examine distinct neuronal populations. Numerous studies have supported the therapeutic approach of targeting tau in AD [54]. The benefits of this approach for LBD are less clear, due to a poor understanding of how tau contributes to LBD pathobiology. Moreso, there is a scarcity of models that emphasize forebrain α-synuclein pathology and neurodegeneration, as limbic and neocortical regions may be more pertinent to comorbid α-synucleinopathy and tauopathy in relation to LBD cognitive dysfunction.

The frequent comorbidity of Lewy pathology with other neuropathologic changes, such as ADNC, LATE-NC, small vessel disease, or granulovacuolar degeneration raises the question of whether these changes co-evolve independently or whether the presence of one pathology influences the initiation and/or maturation of others. An obvious factor to consider is the selective regional basis for some of these neuropathologic changes, where an underlying physiologic vulnerability may be present. In the case of LBD, previous studies have identified the hippocampal CA1 to be related to cognitive dysfunction, with α-synuclein pathology in this subfield correlating with antemortem memory assessment [38]. Intriguingly, the CA1 subfield is also vulnerable to neurofibrillary tangle pathology, to varying levels across neurodegenerative disease [23]. GVBs are frequently detected in CA1 neurons that display early pre-tangle-like alterations to tau and are rarely detected in mature tangle-bearing neurons [47,48]. Thus, GVBs have been implicated as an early tau-related pathological change, but the directionality in which these changes occur is poorly understood, due to the lack of sufficient GVB experimental models. The first mouse model of granulovacuolar degeneration instigated by tau pathology has only recently been developed [31]. Our results demonstrate that α-synuclein pathology may also be sufficient to trigger GVB formation in the mouse hippocampus with substantial enrichment in the CA1 subfield. This is significant given the long-standing association between tau pathology and granulovacuolar degeneration in the human brain. We believe that this mouse model, utilizing α-synuclein PFFs to induce GVB formation in the forebrain, will provide additional clarity in the relationship between α-synuclein and tau aggregation, granulovacuolar degeneration, and, ultimately, neurodegenerative changes.

The causal relationship between α-synuclein, GVB formation, and tau accumulation in our model remains to be further elucidated (**Figure 10**). Given the overlapping pattern of tau puncta and GVB markers within inclusion-bearing neurons, it appears likely that some relationship may be present. Subsequent studies will be useful in identifying the basis for this confluence, whether it be by direct interaction or via dysfunction of endolysosomal pathways. Intriguingly, a recent study identified an interaction between α-synuclein and CHMP2b in a cellular α-synuclein toxicity screen, and that disrupting this interaction was neuroprotective in a mouse α-synucleinopathy model [55]. It is unclear whether a direct interaction with CHMP2b, related to impairments in endosomal sorting complexes required for transport (ESCRT) pathways, might also impact tau independent of GVB formation. ESCRT component knockdown has been shown to promote the propagation of aggregated tau, related to endolysosomal escape [56]. It is possible that the development of α-synuclein pathology in our model engages ESCRT components in parallel with GVB formation, subsequently impacting tau as a result. Beyond this, the nature of tau and GVB accumulation secondary to α-synuclein pathology requires further experimentation. The elucidation of these processes underlying the variable neuronal response to α-synuclein pathology may prove beneficial in furthering our understanding of LBD-related cognitive dysfunction.

**Figure 10.**
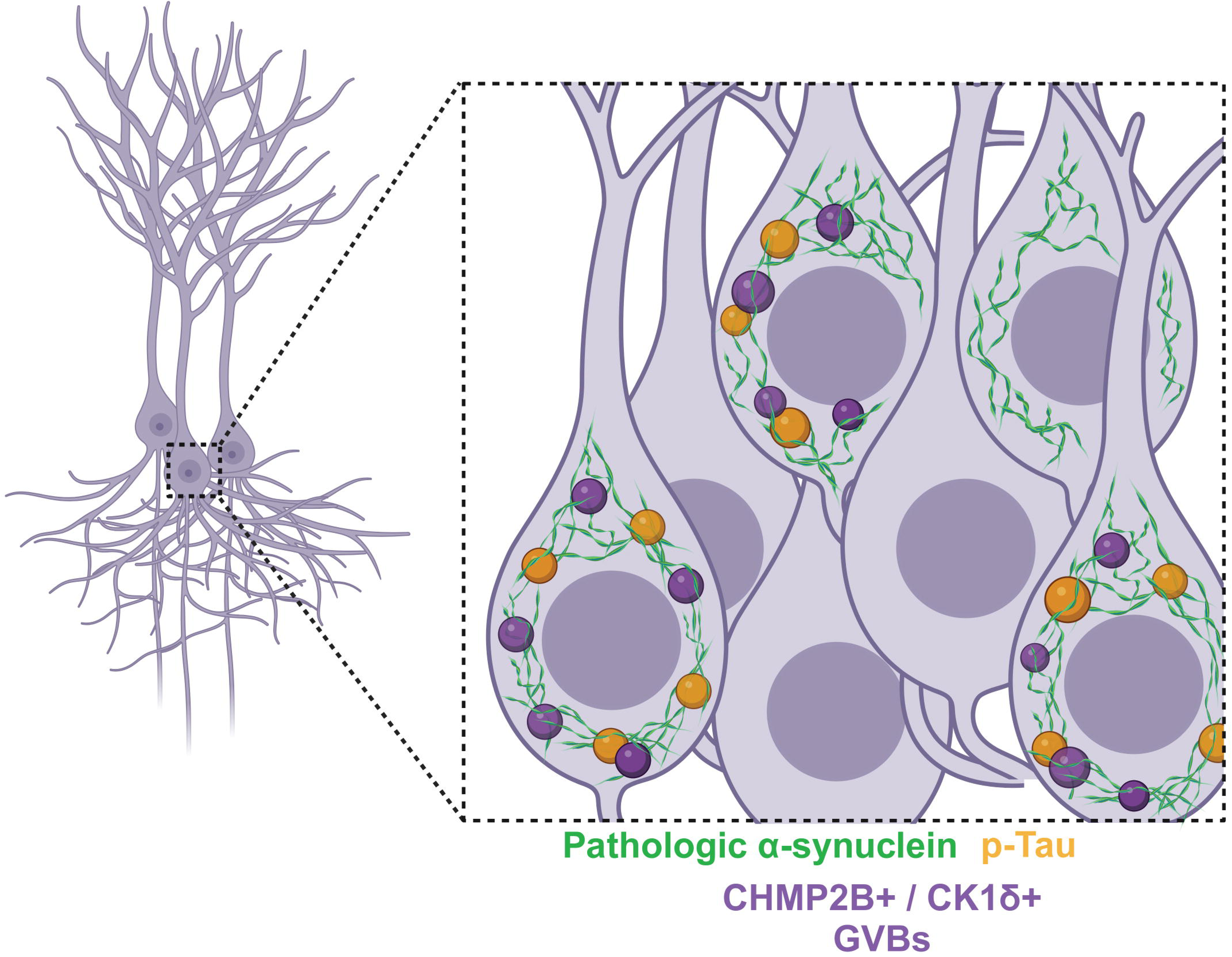
Graphical summary. Image depicts pyramidal neurons harboring a pathologic α-synuclein inclusion accompanied by overlapping phospho-tau and markers of granulovacuolar degeneration bodies (GVBs). GVBs often contain p-tau.

## Conclusions

Stereotactic delivery of α-synuclein PFFs to the basal forebrain of wild-type mice elicits aggregation of pathological α-synuclein throughout the limbic system after 3 months. In this rodent model of LBD, α-synuclein aggregation is accompanied by formation of hyperphosphorylated tau granules and GVBs in many, but not all neuronal subtypes. Notably, neuronal subtypes that form tau granules frequently accumulate GVBs suggesting parallel neuropathological mechanisms between tauopathy and granulovacuolar degeneration. Additionally, the ability of pathological α-synuclein to drive GVB formation without tau suggests a broader relevance of granulovacuolar degeneration to the pathobiology of LBD.

## Methods

### Animals

Wild-type C57BL/6J mice of both male and female sex were obtained from the internal colony of the VAI Vivarium at 10 to 12 weeks of age. *MAPT* knockout mice (B6.129X1-*Mapt^tm1Hnd^*/J) were purchased from Jackson Labs and have been described previously [57]. Housing was limited to 4 mice per cage with food and water provided *ad libitum* under a 12-hour light/dark cycle. Mice were maintained in accordance with NIH guidelines. All studies using mice were reviewed and approved by the Institutional Animal Care and Use Committee (IACUC) of Van Andel Institute.

### Preparation of α-Synuclein PFFs and Stereotactic Surgical Procedures

Mouse α-synuclein pre-formed fibrils (PFFs) were prepared from wild-type, full-length α-synuclein monomers following previously reported methods [58–60]. On the day of surgical procedures, α-synuclein PFFs were thawed to room temperature and pulse sonicated using a Pico 2 sonicator (Biorupter).

Adult C57BL/6J mice were subjected to stereotactic surgery for the delivery of either α-synuclein PFFs or vehicle (PBS). Anesthetized mice were positioned within a Kopf stereotactic frame and the cranium was exposed. For intra-septal (basal forebrain) injection, α-synuclein PFFs (5 µg/µl; 1 µl per site) were delivered in a medial 2-point injection in the basal forebrain using the following coordinates relative to bregma: anterior-posterior (A-P), +0.9 mm; medio-lateral (M-L), 0.0 mm; dorso-ventral (D-V), -3.6 and -4.1 mm. For experiments utilizing intra-hippocampal injection, a lateral 2-point injection into the dorsal hippocampus and overlaying cortex used the following coordinates relative to bregma: anterior-posterior (A-P), -2.5 mm; medio-lateral (M-L), +2.0 mm; dorso-ventral (D-V), -2.4 and -1.4 mm. All injections were delivered in a volume of 1 μl at a flow rate of 0.2 μl/min via blunt tip steel needles (34 gauge) connected through polyethylene tubing to a Hamilton syringe and automatic pump (CMA Microdialysis). Following completion of surgical procedures, mice were removed from anesthesia and monitored until recovery.

### Immunohistochemistry

Mice were perfusion-fixed with 4% paraformaldehyde (PFA) in 0.1 M phosphate buffer (pH 7.4). Post-fixation occurred for 24 hours in 4% PFA followed by cryopreservation in 30% sucrose solution. Whole brains were subsequently processed by microtome into 35 μm-thick coronal sections.

For DAB immunostaining, sections were quenched for endogenous peroxidase activity by incubation in 3% H_2_O_2_ (Sigma) diluted in methanol for 5 min at 4°C. Sections were blocked in 10% normal goat serum (Invitrogen), 0.1% Triton-X100 in PBS for 1 h at room temperature. Sections were incubated with primary antibodies for 48 h at 4°C followed by incubation with biotinylated secondary antibodies (Vector Labs) for 24 h at 4°C. After incubation with ABC reagent (Vector Labs) for 1 h at room temperature and visualization in 3, 3’-diaminobenzidine tetrahydrochloride (DAB; Vector Labs), sections were mounted on Superfrost plus slides (Fisher Scientific), dehydrated with increasing ethanol concentrations and xylene, and coverslipped using Entellan mounting medium (Merck). The following primary antibodies were used: rabbit anti-pS129-α-synuclein (ab51253; Abcam), and mouse anti-pS202/T205-tau (AT8, MN1020, Thermofisher). The following biotinylated secondary antibodies were used: goat anti-mouse IgG and goat anti-rabbit IgG (Vector Labs).

For Gallyas silver staining, sections were fixed in 0.1 M phosphate buffer (pH 7.4) containing 4% PFA for 1 week at 4°C. Sections were then processed with FD NeuroSilver Kit II (FD Neurotechnologies) according to the manufacturer’s instructions. Sections were then mounted on slides, cleared in xylene, and coverslipped with Entellan mounting medium (Merck).

For fluorescence immunostaining, sections were washed in PBS containing 0.1% Triton-X100 then blocked for non-specific binding in PBS containing 10% normal goat serum (Invitrogen) along with 0.4% BSA and 0.2% Triton-X100 for 1 h at room temperature. Sections were incubated in primary antibodies for 48 h at 4 °C and secondary antibodies conjugated to goat anti-rabbit AlexaFluor-488, goat anti-mouse IgG2a AlexaFluor-546, goat anti-guinea pig AlexaFluor-546, goat anti-mouse IgG1 AlexaFluor-647 or goat-anti rat AlexaFluor-647 (Invitrogen) for 2 h at room temperature. The following primary antibodies were used: mouse anti-pS129-α-synuclein (ab184674; Abcam), mouse anti-pS202/T205-tau (AT8, MN1020, Thermofisher), rabbit anti-CHMP2b (ab33174; Abcam), rabbit anti-CK1δ (PA5-32129; Invitrogen), rat anti-LAMP2 (ab13524; Abcam), guinea pig anti-p62 (GP62-C; Progen), rabbit anti-TFE3 (ab93808; Abcam). Sections were then washed in PBS + 0.1% Triton, followed by a 30 min incubation in PBS containing DAPI. Sections were subsequently mounted on Superfrost plus slides (Fisher Scientific) and coverslipped using Prolong mounting medium (Invitrogen).

### Digital Slide-Scanning and Quantitative Pathology

Slide-mounted coronal brain sections were scanned using a ScanScope XT slide scanner (Aperio) at a resolution of 0.5 µm/pixel. Annotation and quantitative assessment of the hippocampal subfields across the anterior-posterior axis (6 coronal sections assessed per animal, left/right hemispheric regions were annotated and assessed jointly) were performed using HALO analysis software (Area quantification, Object colocalization; Indica Labs Inc.). For each quantitative approach, threshold parameters were developed and optimized to detect positive-staining without background staining, tissue-level artifacts, or edge-effects. All sections/slides were assessed using the optimized HALO batch analysis function for each marker of interest.

### Confocal Imaging of Fluorescence Immunostaining

Confocal images were acquired in x, y, and z planes using an ImageXpress Confocal HT.ai high-content microscope (Molecular Devices) equipped with a 60x water-immersion objective and a 60-micron pinhole spinning disc. Maximum intensity projections were used for image analysis. Z-steps information can be found in **Table S1**.

### Fluorescence Immunohistochemistry Image Analysis

#### Quantification of Granulovacuolar Degeneration Bodies in the Distal CA1 Subfield

Maximum intensity projections of 60X tiled and stitched images were analyzed using the granularity module in the MetaXpress analysis software (Molecular Devices). The number of puncta per area and percent area occupied of CHMP2b or CK1δ as well as pS202/T205-tau and pS129-α-synculein were measured in the CA1 subfield of the hippocampus. 2 images per mouse were analyzed from 2-3 Sham or PFF treated mice. Each data point represents measurements from individual tiled and stitched images corresponding to the CA1 subfield of the hippocampus from distinct hemispheres.

#### Quantification of Enlarged Lysosomes in the CA3

Enlarged lysosomes were identified and quantified using NIH ImageJ (FIJI). Bright LAMP2-positive lysosomes were identified by adjusting image thresholds to 3000, 65535. Particles smaller than 150 pixels were removed before running particle analysis to measure enlarged LAMP2 particles. Total particle area, number of particles and average particle size were recorded. 3-4 images were analyzed per mouse. Data is shown as the mean for each measurement per mouse.

### Primary Neuronal Culture

Primary neuronal cultures were prepared from embryonic day 18 (E18) CD1 mice. Brains were gently removed from the embryos and placed into a petri dish filled with ice-cold, sterile Hibernate Medium (Cat#. A1247601 Gibco). The hemispheres were gently separated, and the meninges, thalamus, striatum, brainstem, and hippocampus were removed. Hippocampal tissue isolated from each hemisphere was pooled and digested in papain solution (20 U/mL Cat# LS003126, Worthington) and then treated with DNase I (Cat# LS006563, Worthington) to remove residual DNA. The tissue was then washed with pre-warmed Neurobasal media (Cat# 21103049 Gibco), mechanically dissociated, and strained through a 40 μm cell strainer. The cell suspension was pelleted at 1000 rpm for 5 min, resuspended in 2 mL of neuron media (Neurobasal media containing 1% B27, 2 mM GlutaMAX, and penicillin-streptomycin), and gently mixed. The dissociated neurons were seeded on poly-*D*-lysine (Cat# P0899, Sigma) coated 96-well culture plates (Cat# 6005182, Perkin Elmer) at 17,000 cells/well. Cells were maintained at 37°C in 5% CO_2_.

### Immunocytochemistry and Imaging Analysis

Cells were fixed with prewarmed 4% PFA containing 4% sucrose for 15 mins, followed by 5 washes with PBS. Cells were then permeabilized with 0.3% Triton X-100 in 3% bovine serum albumin (BSA) for 15 mins followed by a 5X wash with PBS. Cells were blocked with 3% BSA for 1 h before overnight incubation with primary antibodies at 4°C with shaking. Cells were then incubated with respective AlexaFluor secondary antibodies for 1 h at room temperature with shaking, followed by 5X wash with PBS. Cells were then stained with DAPI (0.1 ug/mL), and plates were sealed and stored at 4°C. Plates were imaged at 20x magnification with a 0.5X digital zoom ten ROIs per well using a Zeiss Cell Discoverer 7 microscope. Images were processed in Zen 3.0 software. Cells were detected on DAPI signal with an expansion ratio of 9 to encase the soma of the neurons. Cells were then classified as being positive for one or both markers used in the ICC based on threshold ratios of true positives to background signal. The total cell and double-positive cell count of each ROI were averaged within the same well. The ratio of double-positive cells to total cells was then normalized to the 0.2 μg/mL dose of α-synuclein PFFs. Data points were derived from 1-2 wells per plate from 4 separate cultures.

### Behavior

A dedicated mouse behavioral testing suite was used for all behavioral assessments. ANY-maze software (Stoelting) was used in conjunction with a video-tracking system to assess behavior during all testing. During testing, mice were randomly intermixed during testing in a blinded manner regarding treatment group. Prior to testing, mice were habituated to handling and allowed to acclimate to the behavioral testing suite. A minimum rest period of 24 hours was provided between assays that did not require multiple sequential days of testing. Behavioral apparatus used in testing was thoroughly cleaned with 70% ethanol between trials.

#### Elevated-Plus Maze

Mice were initially placed in an enclosed corner of the maze apparatus and monitored via over-head placement of a video camera, with the examiner hidden from view of the open arms of the apparatus. Over a single testing period of 10 mins, movement was tracked and the number of open versus closed arm entries (defined as full body entry based on body detection) as well as time spent in each arm was determined. Between assessments, the apparatus platform was cleaned with 70% ethanol.

#### Y-Maze

Mice were initially place in the center of the maze apparatus and monitored with over-head placement of a video camera. Over a testing period of 10 mins, the number of entries (defined as full body entry based on body detection) into each of the three corridors of the Y-maze apparatus were determined. A calculation determining the % spontaneous alternation rate was then performed to determine the number of non-repeat corridor entries relative to total entries. Between assessments, the Y-maze apparatus was cleaned with 70% ethanol.

#### Barnes Maze

A white Barnes maze platform with equi-distant spacing of escape holes along the outer perimeter was used for all trials, with a single hole replaced with a larger escape pod. An acquisition phase of 3 trials per day was followed for a total of 4 days (12 total trials per animal). The maximum trial duration was limited to 3 mins with a 20-min inter-trial interval. The duration for the animals’ head to enter the target hole (latency) during each trial was evaluated using ANY-maze software with a module specifically developed for the assay. Following completion of the acquisition phase, a probe phase occurred 24 hours later. During the probe phase, the duration of time spent in the quadrant previously holding the escape hole was determined. Between trials, the Barnes maze platform was cleaned with 70% ethanol.

#### Fear Conditioning

To assess fear learning and memory, mice were placed in an isolation cubicle within a larger fear conditioning system (Ugo Basile). During a period of fear acquisition, mice were trained to associate a tone with mild foot shock (0.8 mA) that lasted for 2-seconds. Following a 2-minute period, the tone would play for 30 s followed by a mild foot shock. This was repeated several times to facilitate fear learning. A fear response (freezing) was identified via video-tracking system coupled with ANY-maze software (Stoelting). To assess contextual fear conditioning, mice were placed in the isolation cubicle with the same visual cues and fear response was evaluated over a 5-minute testing period. After a 2-hour recovery period, mice were then placed in the isolation cubicle with novel visual cues and fear response was again evaluated over a 5-minute period. To assess cued fear conditioning, the tone was played for 30 s without any foot shock. The tone was repeated after a 2-minute period. Fear response during each of the tones was evaluated.

### Human Tissue Immunostaining

#### Case Information

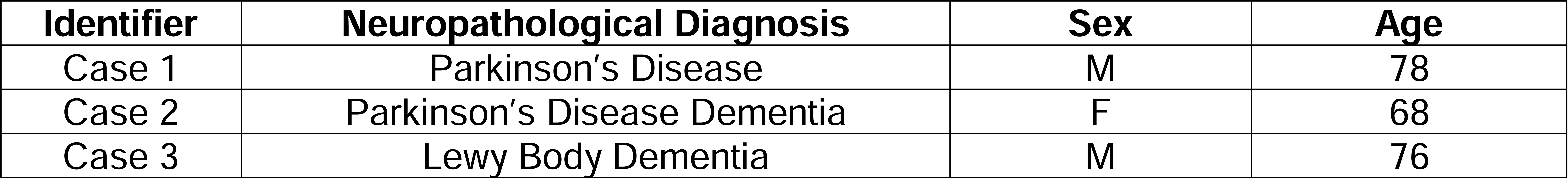

#### Immunohistochemistry

For pS129-α-synuclein, slides were deparaffinized in xylene and rehydrated in a descending ethanol series and then rinsed in DI water for 10 mins. Slides were then incubated in antigen unmasking solution (H-3300, Vector Labs) and microwaved for 15 mins at 95°C. Slides were allowed to cool for 20 mins at room temperature and washed in running tap water for 10 mins. Slides were then incubated in 5% H_2_O_2_ in water, or 30% H_2_O_2_ in methanol, to quench endogenous peroxidase activity for 30 mins at room temperature, followed by 10 mins of running tap water and then 5 mins in 0.1 M Tris Buffer pH 7.6. Slides were then blocked in 0.1 M Tris/2% fetal bovine serum (FBS) for 1 hour followed by incubation with primary antibody diluted in 0.1 M Tris/2% FBS overnight at 4°C. Primary antibodies include: rabbit anti-pS129-α-synuclein antibody (ab51253, Abcam) 1:20,000, rabbit anti-CHMP2b (ab33174; Abcam) 1:250 and rabbit anti-CK1δ (PA5-32129; Invitrogen) 1:250. After primary antibody incubation, slides were rinsed with 0.1 M Tris for 5 mins, then incubated with goat anti-rabbit biotinylated IgG (BA1000, Vector Labs) in 0.1 M Tris/2% FBS 1:1000 for 1 hour. Biotinylated antibody was rinsed off with 0.1 M Tris for 5 mins, then incubated with avidin-biotin complex solution (PK-6100, Vector Labs) for 1 hour. Slides were washed in 0.1M Tris. Slides were developed in DAB solution (ImmPACT DAB Peroxidase Substrate, SK-4105, Vector Labs). After DAB development, slides were washed in diH_2_O and briefly counterstained in Harris Hematoxylin (6765001, Thermo Fisher) or Dako EnVision Flex hemotoxylin (K800821-2, Dako). Slides were dehydrated in an ascending ethanol series and cleared in xylene before cover slipping with Cytoseal 60 (22-244-256, Fisher Scientific).

For pSer202/Thr205-tau, slides were deparaffinized and retrieved with a Dako PT link, using Dako Env FLEX High pH buffer pH 8.8 (K800421-5, Dako) for 20 mins at 97°C. Post retrieval, slides were transferred to a Dako Autostainer link 48 where the remaining staining steps were performed on the automated platform, utilizing Dako EnV FLEX, High pH kit reagents (K800021-5, Dako). pSer202/Thr205-tau antibody (MN1020B, Invitrogen) was diluted to 1:2000 in Dako Env FLEX Ab diluent (K800621-2, Dako) for 30 mins, followed by 5 min incubation with Dako wash buffer. Slides were incubated with Dako Env FLEX/HRP polymer for 20 mins followed by a 5 min rinse in Dako wash buffer. Chromogen development was performed using a 10 min incubation followed by 5 min water rinse and a 10 min counterstain with Dako EnVision Flex hemotoxylin (K800821-2, Dako). Slides were transferred to Sakura Prisma Autostainer for dehydration and coverslipping.

#### Immunofluorescence

6 μm paraffin-embedded tissue sections were acquired through the Van Andel Institute Brain Bank. Slides were de-paraffinized with 2 sequential 5-min washes in xylene and then were rehydrated in 1-min washes in a descending series of ethanol: 100%, 100%, 95%, 80%, 70%. Slides were washed in deionized water for 1 min prior to transfer to the BioGenex EZ-Retriever System where they were incubated in an antigen unmasking solution (Vector Laboratories; Cat# H-3300) at 95°C for 15 min. Slides were left to cool at room temperature for 20 mins and then washed in running tap water for 10 mins. Slides were washed for 5 mins in 0.1 M Tris prior to blocking for 1 hour in 0.1 M Tris/2% fetal bovine serum (FBS). Slides were incubated in primary antibody in 0.1 M Tris/2% FBS in a humidified chamber overnight at 4°C; pS129-α-synuclein (81A), (825701, BioLegend, 1:1000); phospho-tau (Ser202, Thr205) (AT8), (MN1020, Thermo Fisher 1:1000); CK1δ, (PA5-32129 Thermo Fisher, 1:250); CHMP2b, (ab33174 Abcam 1:250). The primary antibodies were rinsed off in 0.1 M Tris for 5 mins and incubated with secondary antibodies goat anti-mouse IgG2a AlexaFluor-488, goat anti-rabbit IgG AlexaFluor-568, goat anti-mouse IgG1 AlexaFluor-647 (Thermo Fisher) on slides in the dark for 2 hours at room temperature. Slides were washed in 0.1 M Tris for 5 mins and then immersed in Sudan Black (0.3% Sudan Black in 70% ethanol) for 15 secs to quench autofluorescence. Slides were washed for 5 mins three times in 0.1 M Tris in the dark. Coverglass was then mounted on slides with ProLong gold with DAPI (Invitrogen, Cat#P36931). Slides were imaged on an ImageXpress Confocal HT.ai High-Content Imaging System at 60x magnification.

## Abbreviations

LBD: Lewy Body Dementia
GVBs: granulovacuolar degeneration bodies
PFFs: pre-formed fibrils
ADNC: Alzheimer’s disease neuropathologic change
PD: Parkinson’s disease
PDD: Parkinson’s disease dementia
DLB: Dementia with Lewy bodies
AD: Alzheimer’s disease

## Data Availability

All data generated or analyzed during this study are included in this published article and its supplementary information.

## Ethical approval and consent to participate

All studies using mice were reviewed and approved by the Institutional Animal Care and Use Committee (IACUC) of Van Andel Institute. All studies involving human brain samples were reviewed and approved by the Institutional Review Board of Van Andel Institute. Informed consent for brain donation was obtained by the Van Andel Institute’s organ procurement organization partners and all donors remain anonymous.

## Consent for publication

All authors read and approved the final manuscript.

## Author Contributions

DJD and DJM conceptualized this research. DJD and APT performed stereotactic surgeries. DJD, MLE, AK, NM and ETW performed and analyzed experiments. MXH and DJM supervised this research. DJD, MLE and DJM wrote the manuscript.

## Acknowledgements

We would like to thank the patients and families who participated in this research, without whom this study would not have been possible. Human brain tissue was provided by the Van Andel Institute Brain Bank (RRID:SCR_026035), which is supported by the West Michigan Neurodegenerative Diseases (MiND) Program. We also thank the Van Andel Institute Vivarium (RRID:SCR_023211), Optical Imaging (RRID:SCR_021968), and Pathology and Biorepository (RRID:SCR_022912) Cores for technical assistance. We would also like to thank the MiND iPSC & Screening Platform. Some figures were created using Biorender (BioRender.com).

