## Supplemental Figures for "Pathological α-synuclein elicits granulovacuolar degeneration independent of tau"

### Supplemental Materials

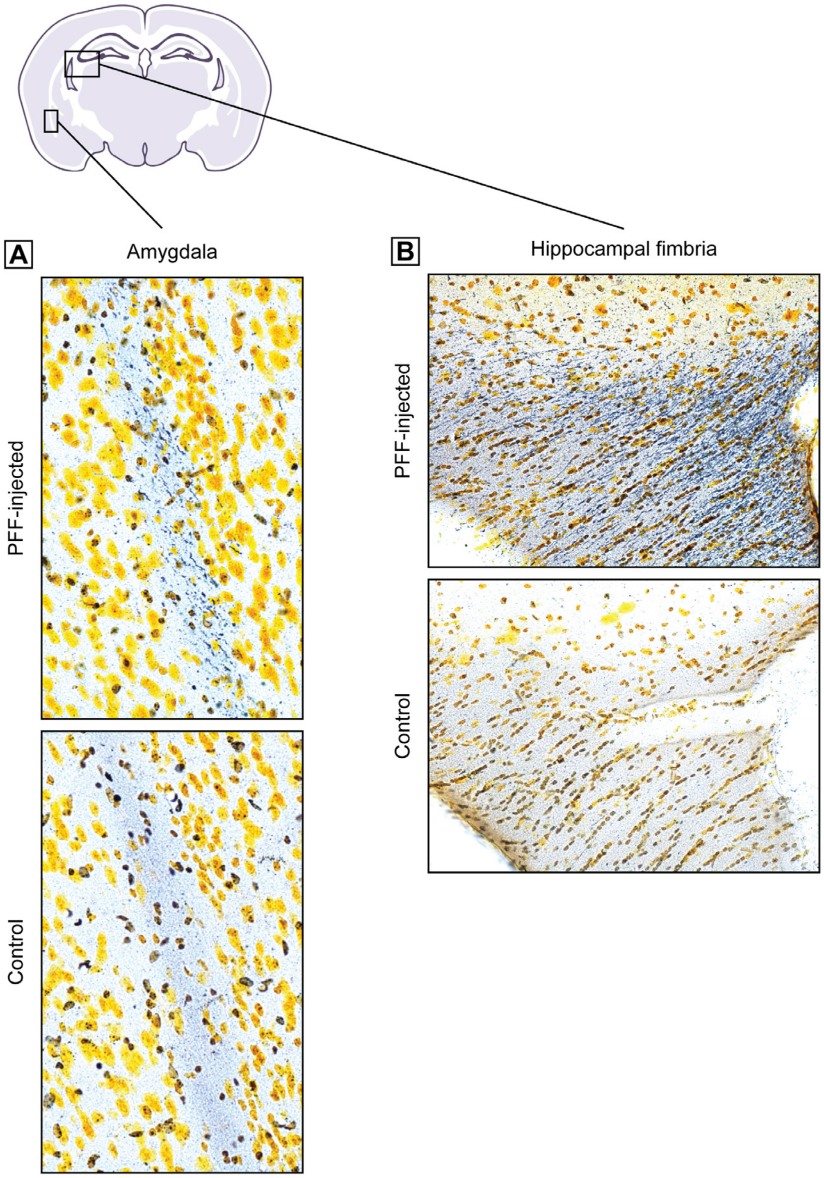


**Figure S1. The basal forebrain PFF-injection paradigm elicits minimal neurodegenerative changes at 3 MPI. A-B)** Representative histological sections depicting modest Gallyas silver-positive neuronal processes (black fibers) in the **(A)** amygdala capsule and **(B)** hippocampal fimbria at 3 MPI.

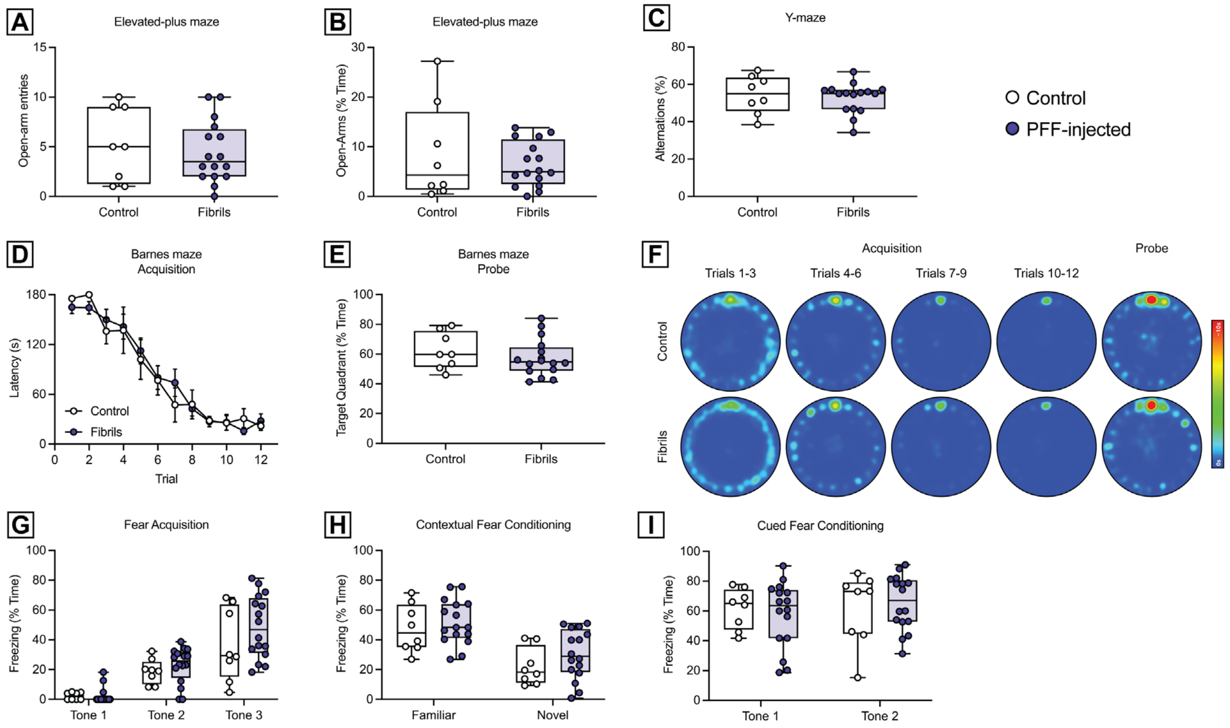


**Figure S2. Basal forebrain PFF-injected mice do not exhibit cognitive or anxiety-related deficits at 3 MPI. A-B)** Performance in the elevated-plus maze as measured by **(A)** open-arm entries and **(B)** % time spent in open-arms. Data are expressed as boxplots depicting the median, interquartile range, and individual data points (*n* = 8-16 mice/group). Non-significant by unpaired Student's *t*-test. **C)** Performance in the Y-maze as measured by % spontaneous alternations. Data are expressed as boxplots depicting the median, interquartile range, and individual data points (*n* = 8-16 mice/group). Non-significant by unpaired Student's *t*-test. **D-E)** Performance in the Barnes maze **(D)** acquisition as measured by latency in seconds and **(E)** probe sessions measured by % time in the target quadrant. Data are expressed as boxplots depicting the median, interquartile range, and individual data points of all mice (*n* = 8-16 mice/group). Non-significant by two-way ANOVA with Bonferroni's multiple comparisons test or unpaired Student's *t*-test, respectively. **F)** Heatmap depicting the average location of all assessed mice per group in relation to time (trial sessions) in the Barnes maze. **G-I)** Performance in the fear conditioning assay, as measured by % freezing time during **(G)** fear acquisition, **(H)** contextual fear conditioning, and **(I)** cued fear conditioning. Data are expressed as boxplots depicting the median, interquartile range, and individual data points of all mice (*n* = 8-16 mice/group). Non-significant by unpaired Student's *t*-test.

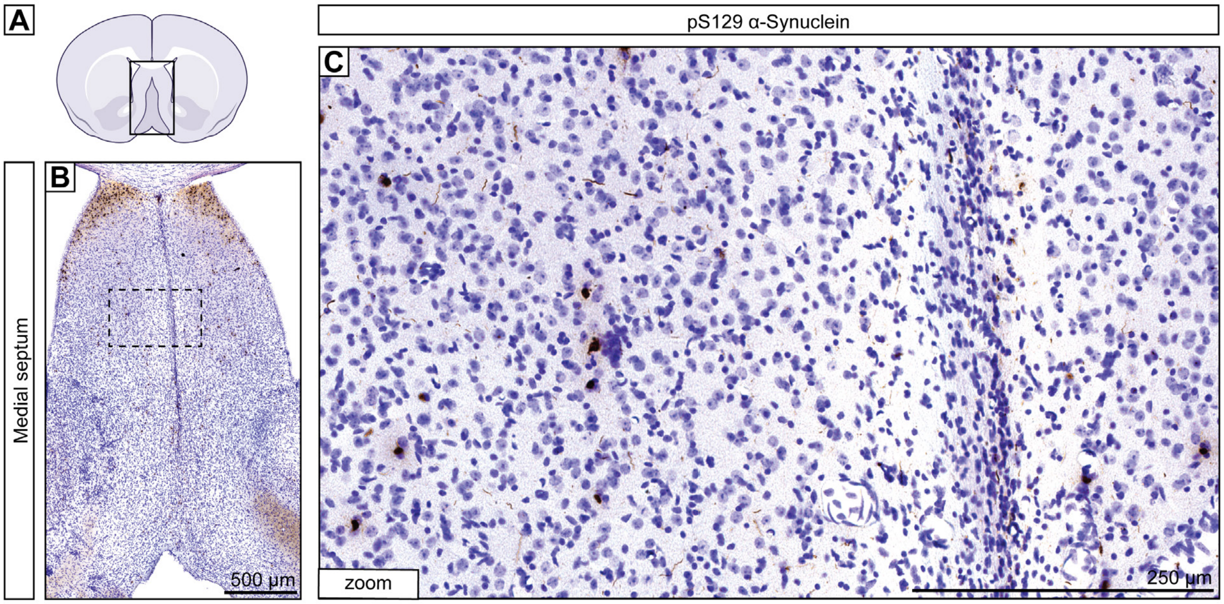


**Figure S3. Minimal pathological α-synuclein inclusions are observed within the medial septal area proximal to the needle track. A)** Representative schematic of a mouse coronal brain section highlighting the location of the basal forebrain. **B-C)** Higher magnification images of the medial septal area depicting pS129-α-synuclein inclusions in relation to the needle track. Scale bar: (**B**) 500 µm, and (**C**) 250 µm.

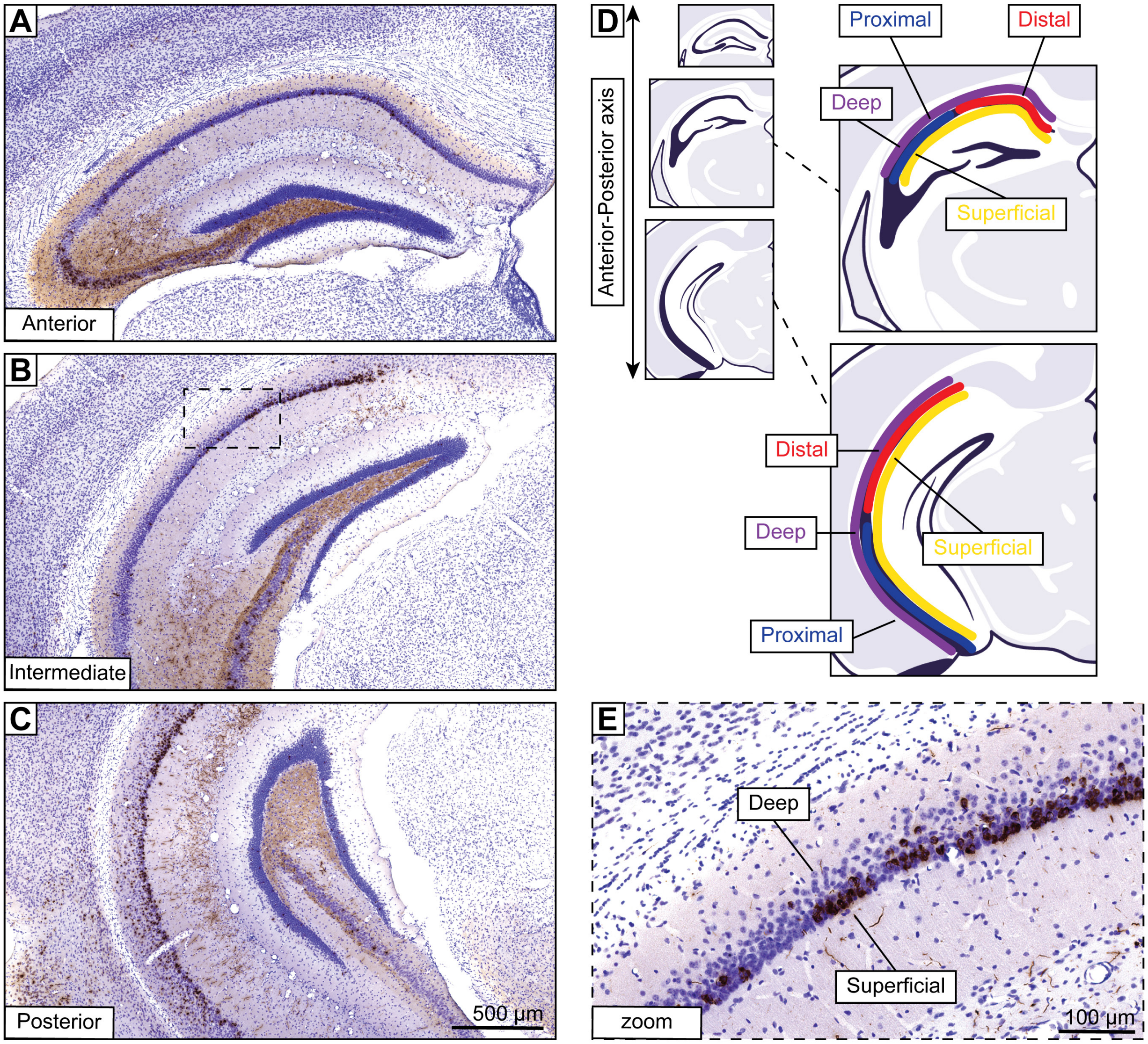


**Figure S4. Distribution of α-synuclein pathology across the hippocampal axis. A-C)** Representative mouse coronal brain sections displaying pS129-α-synuclein immunostaining with Nissl counterstain. The hippocampus is shown along the **(A)** anterior, **(B)** intermediate, and **(C)** posterior portions of the axis. Scale bar: 500 µm. **D)** Representative schematic of the mouse hippocampal anterior-posterior axis with labeling of the distal/proximal portions and deep/superficial layers along the CA1 subfield. **E)** Inset image of panel **(B)** displaying higher magnification view of the distal CA1 pyramidal layer with pS129-α-synuclein inclusions predominantly localized to the superficial layer. Scale bar: 100 μm.

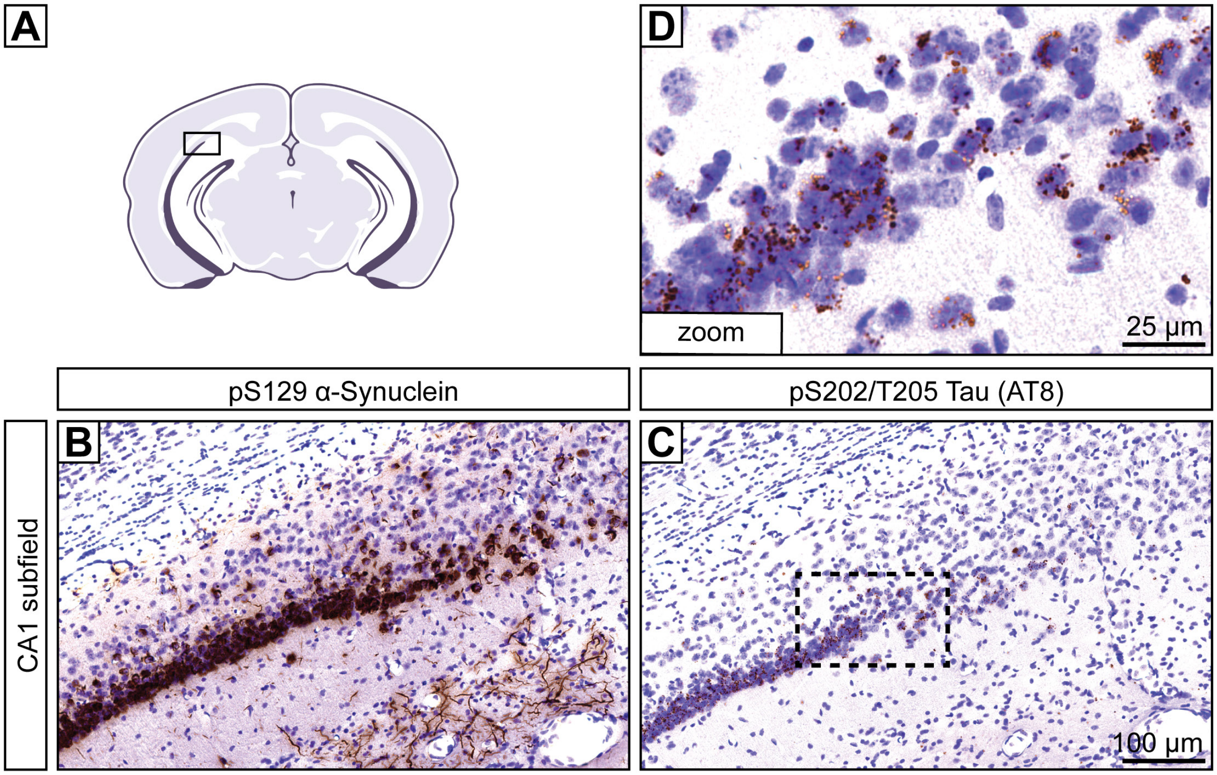


**Figure S5. The distal CA1 pyramidal layer of the intermediate to posterior hippocampus is heavily affected by both α-synuclein and tau pathology. A)** Representative schematic showing the region of interest in a mouse coronal section featuring the posterior hippocampus. **B-C)** Representative histological images displaying either **(B)** pS129-α-synuclein or **(C)** pS202/T205-tau (AT8) immunostaining with Nissl counterstain. Scale bar: 100 µm. **D)** Abundant AT8+ tau puncta are observed in the boxed region from panel **(C)**. Scale bar: 25 µm.

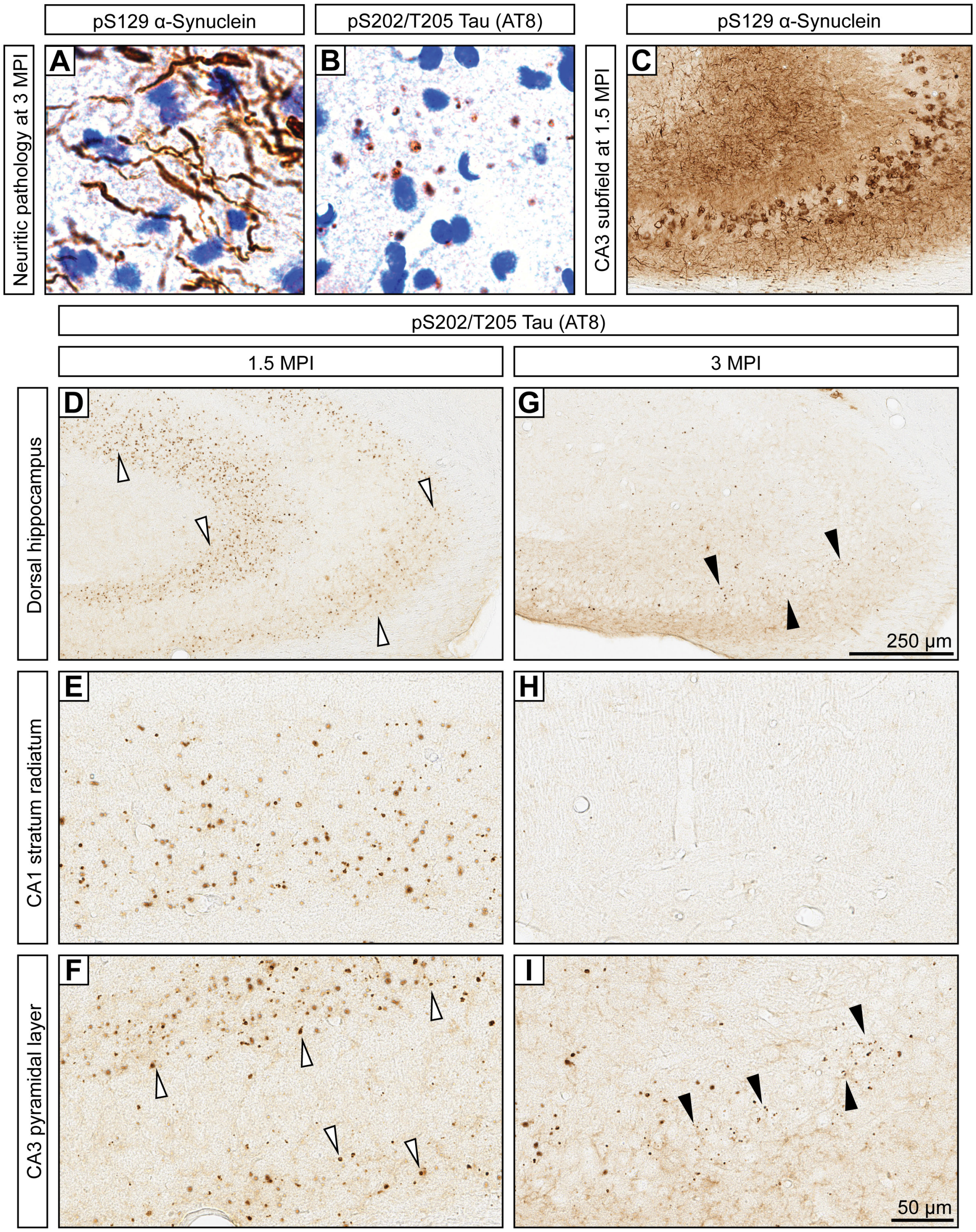


**Figure S6. Prior to 3 MPI, tau puncta are more prominent in neuronal processes of the hippocampus. A)** At 3 MPI, neuritic pS129-α-synuclein pathology is detected in the hippocampus. **B)** Correspondingly, neuritic tau (AT8) puncta are also observed at 3 MPI.

**C)** Basal forebrain injected mice exhibit robust α-synuclein pathology in the hippocampal CA3 subfield at 1.5 MPI. **D-F)** At 1.5 MPI, larger tau puncta are more abundant in the stratum oriens and **(E)** stratum radiatum layers of the hippocampus, within neuronal processes, with fewer small puncta observed in the **(D)** pyramidal layer. **G-I)** At 3 MPI, tau puncta are more abundant in the **(I)** pyramidal layer, in cell bodies, with fewer puncta observed in the **(H)** process layers. Scale bars: **(D, G)** 250 μm and **(E, F, H, I)** 50 μm

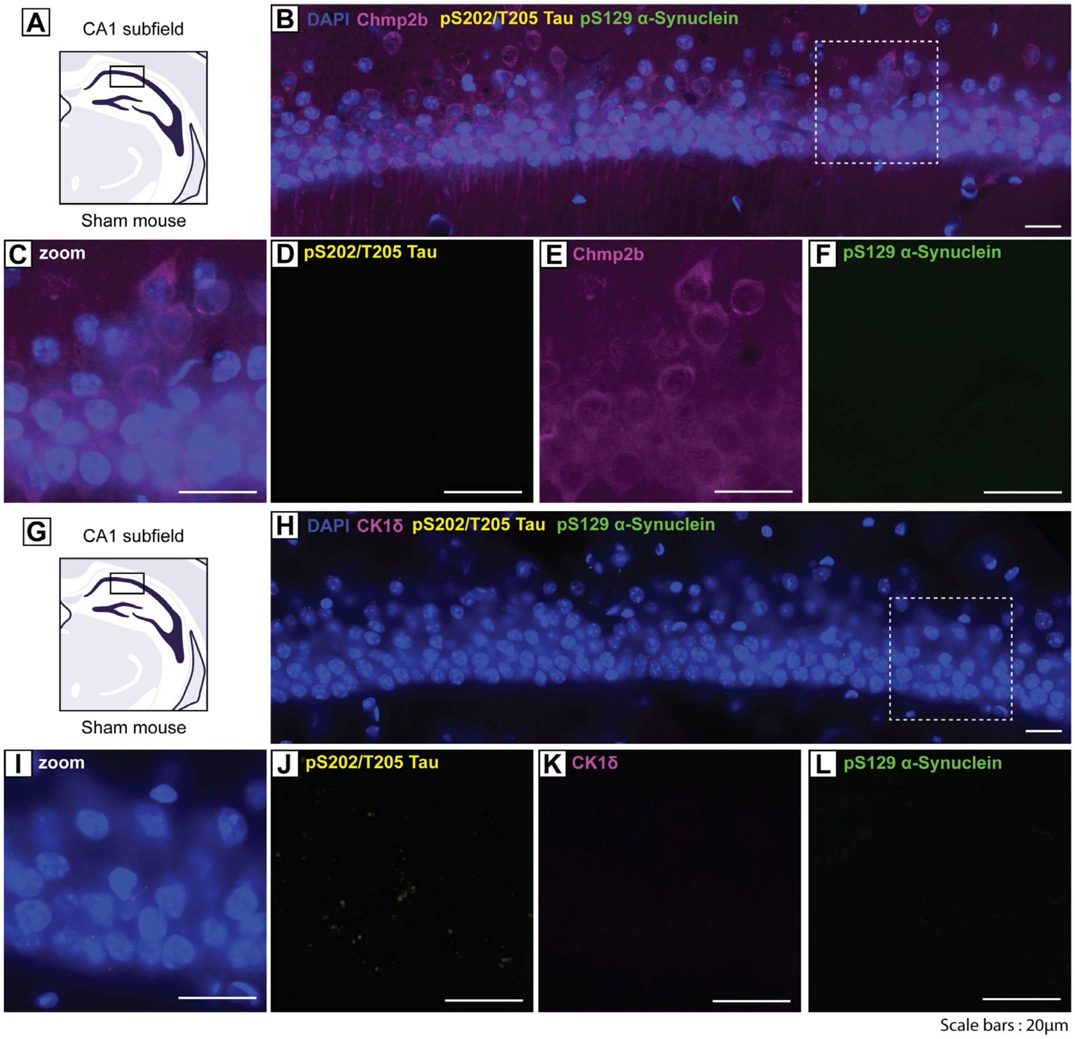


**Figure S7. Absence of α-synuclein, tau, and granulovacuolar degeneration bodies in the distal CA1 subfield of sham-injected control mice. A, G)** Representative schematic of the distal CA1 pyramidal layer of the intermediate/posterior hippocampus of Sham-injected mice. **B)** Confocal immunofluorescent image showing pS129-α-synuclein, pS202/T205-tau, and Chmp2b in the CA1 of sham mice. **C-F)** Inset image from **(B)** showing **(C)** combined, **(D)** pS202/T205-tau, **(E)** Chmp2b, and **(F)** pS129-α-synuclein. **H)** Confocal immunofluorescent image showing pS129-α-synuclein, pS202/T205-tau, and CK1δ in the CA1 of Sham-injected mice. **I-L)** Inset image from **(H)** showing **(I)** combined, **(J)** pS202/T205-tau **(K)** CK1δ, and **(L)** pS129-α-synuclein. Scale bars: 20 μm.

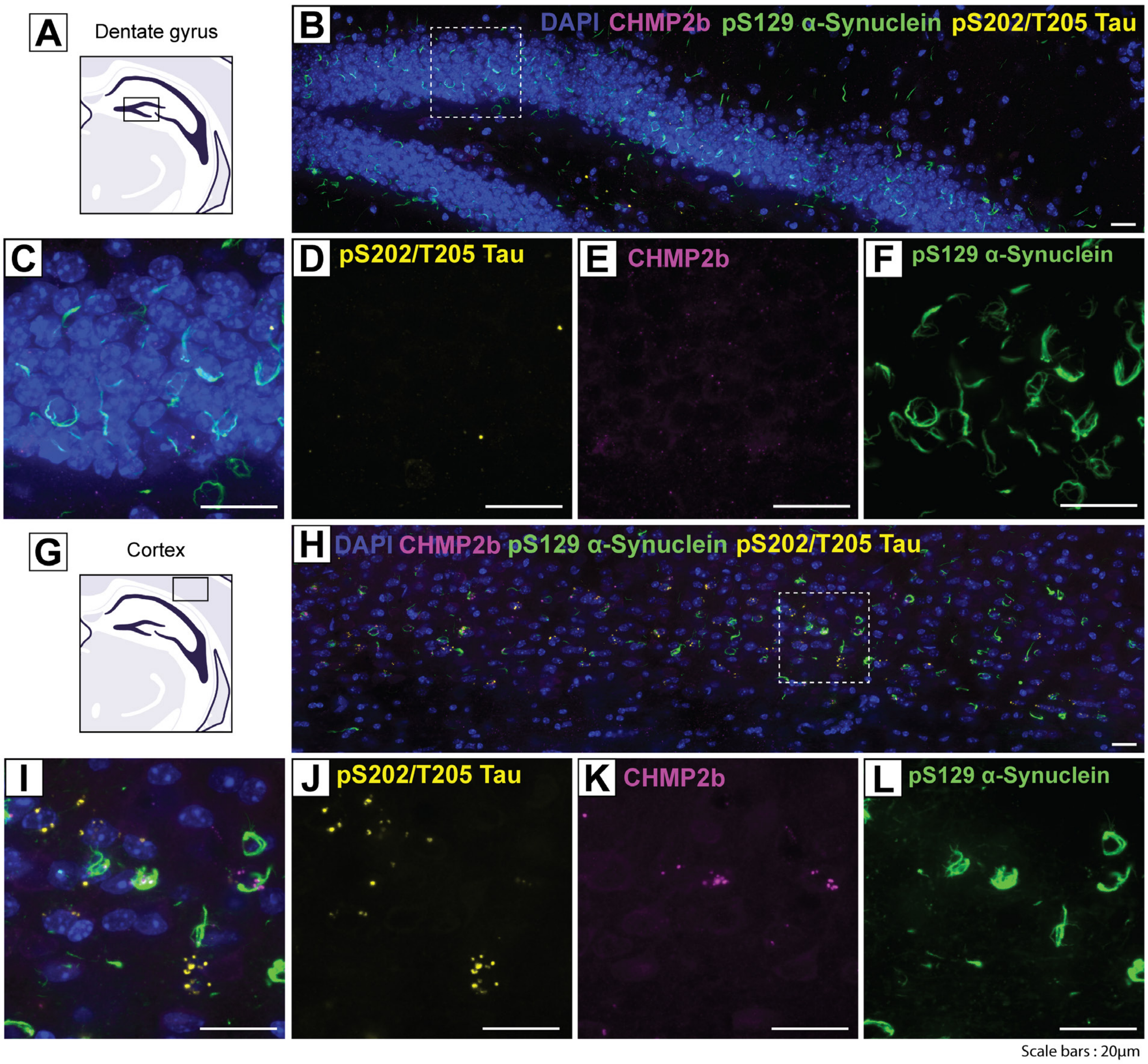


**Figure S8. Cellular localization of α-synuclein, tau, and granulovacuolar degeneration bodies in the hippocampal PFF-injection paradigm. A)** Representative schematic of the dentate gyrus. **B)** Confocal immunofluorescent image showing pS129-α-synuclein, pS202/T205-tau, and Chmp2b in the dentate gyrus. **C-F)** Inset image from **(B)** showing **(C)** combined, **(D)** pS202/T205-tau, **(E)** Chmp2b, and **(F)** pS129-α-synuclein. **G)** Representative schematic of the cortex overlaying the hippocampus. **H)** Confocal immunofluorescent image showing pS129-α-synuclein, pS202/T205-tau, and Chmp2b in the cortex. **I-L)** Inset image from **(H)** showing **(I)** combined, **(J)** pS202/T205-tau **(K)** Chmp2b, and **(L)** pS129-α-synuclein. Scale bars: 20 μm.

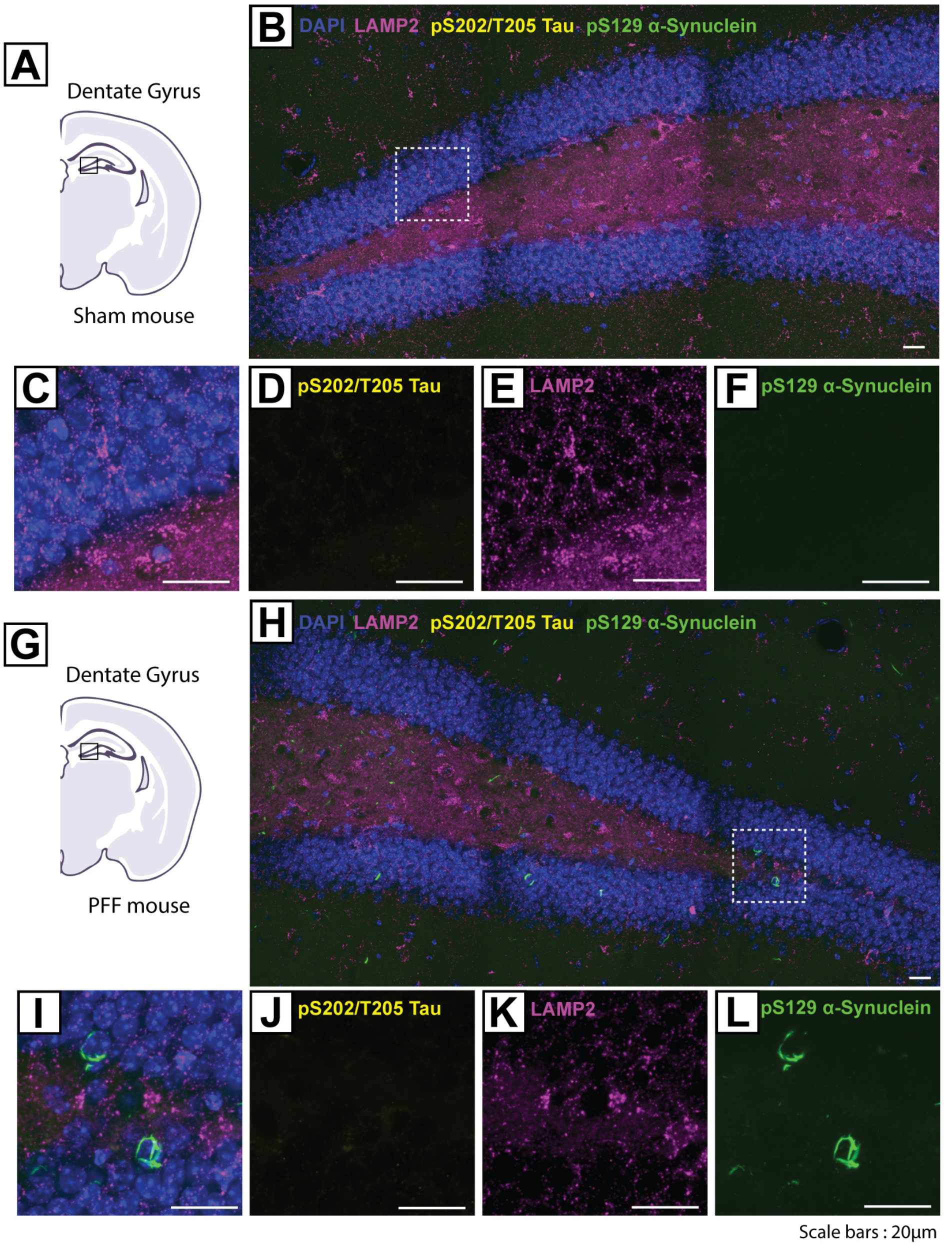


**Figure S9. pS129-α-synuclein accumulates in the granule cell layer of the dentate gyrus without pS202/T205-tau inclusions or enlarged lysosomes A, G)** Representative schematic of the dentate gyrus of the intermediate/posterior hippocampus in **(A)** Sham or **(G)** PFF-injected mice. **B)** Confocal immunofluorescent image showing pS129-α-synuclein, pS202/T205-tau, and LAMP2 in the dentate gyrus of Sham mice. **C-F)** Inset image from **(B)** showing **(C)** combined, **(D)** pS202/T205-tau, **(E)** LAMP2, and **(F)**

pS129-α-synuclein. **H)** Confocal immunofluorescent image showing pS129-α-synuclein, pS202/T205-tau, and LAMP2 in the CA1 of PFF mice. **I-L)** Inset image from **(H)** showing **(I)** combined, **(J)** pS202/T205-tau, **(K)** LAMP2, and **(L)** pS129-α-synuclein. Scale bars: 20 μm.

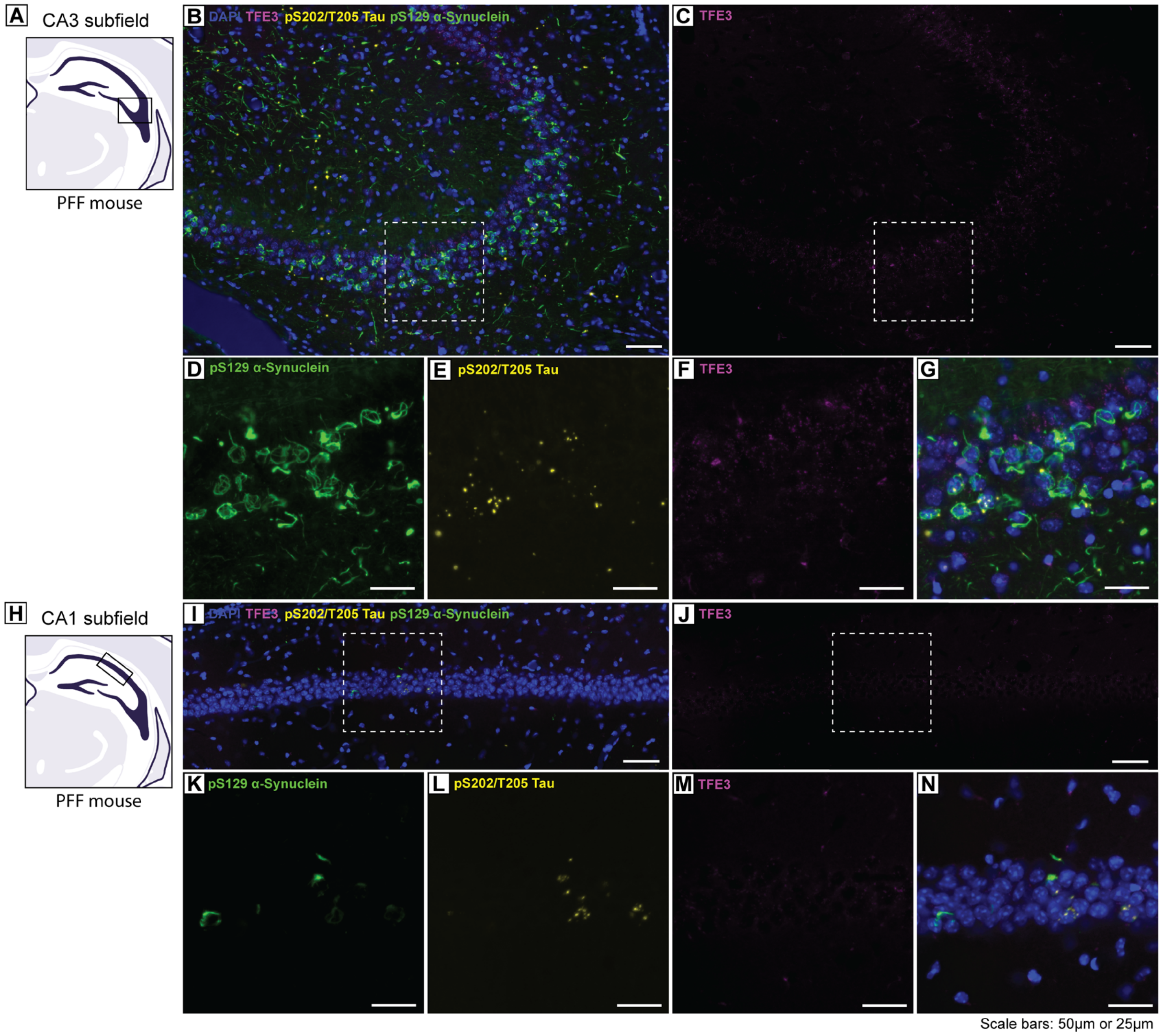


**Figure S10. TFE3 localization in basal forebrain PFF-injected mice. A, H)** Representative schematic of **(A)** the CA3 of the intermediate/posterior hippocampus or **(H)** the distal CA1 pyramidal layer of the intermediate/posterior hippocampus of PFF-injected mice. **B)** Confocal immunofluorescent image showing pS129-α-synuclein, pS202/T205-tau, and TFE3 in the CA3. **C)** TFE3 in the CA3. **D-G)** Inset image from **(B)** showing **(D)** pS129-α-synuclein. **(E)** pS202/T205-tau, **(F)** TFE3, and **(G)** combined. **I)** Confocal immunofluorescent image showing pS129-α-synuclein, pS202/T205-tau, and TFE3 in the CA1 of PFF mice. **J)** TFE3 in the CA1. **K-N)** Inset image from **(I)** showing **(K)**

pS129-α-synuclein. **(L)** pS202/T205-tau, **(M)** TFE3, and **(N)** combined. Note, TFE3 nuclear translocation or altered expression levels are not detected. Scale bars: 50 μm **(B, C, I, J)** or 25 μm **(D-G, K-N).**

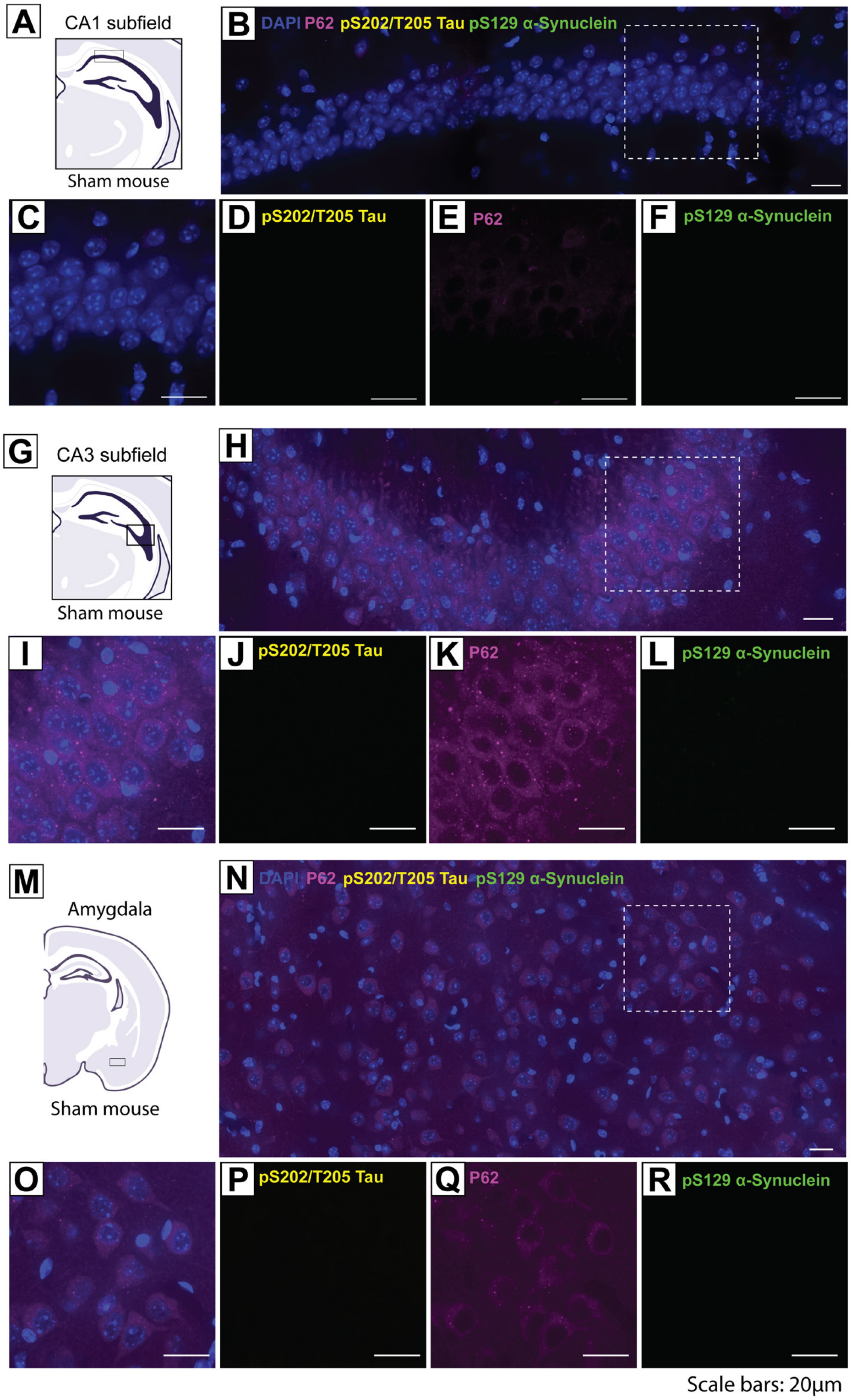


**Figure S11. p62 localization in Sham-injected mice. A, G, M)** Representative schematic of **(A)** the distal CA1 pyramidal layer of the intermediate/posterior hippocampus, **(G)** the CA3 of the intermediate/posterior hippocampus or **(M)** the amygdala in Sham-injected control mice. **B)** Confocal immunofluorescent image showing pS129-α-synuclein, pS202/T205-tau, and p62 in the CA1. **C-F)** Inset image from **(B)** showing **(C)** combined, **(D)** pS202/T205-tau, **(E)** p62, and **(F)** pS129-α-synuclein. **H)** Confocal immunofluorescent image showing pS129-α-synuclein, pS202/T205-tau, and p62 in the CA3. **I-L)** Inset image from **(H)** showing **(I)** combined, **(J)** pS202/T205-tau, **(K)** p62, and **(L)** pS129-α-synuclein. **N)** Confocal immunofluorescent image showing pS129-α-synuclein, pS202/T205-tau, and p62 in the amygdala. **O-R)** Inset image from **(N)** showing **(O)** combined, **(P)** pS202/T205-tau, **(Q)** p62, and **(R)** pS129-α-synuclein. Note, p62 is mostly cytoplasmic. Scale bars: 20 μm.

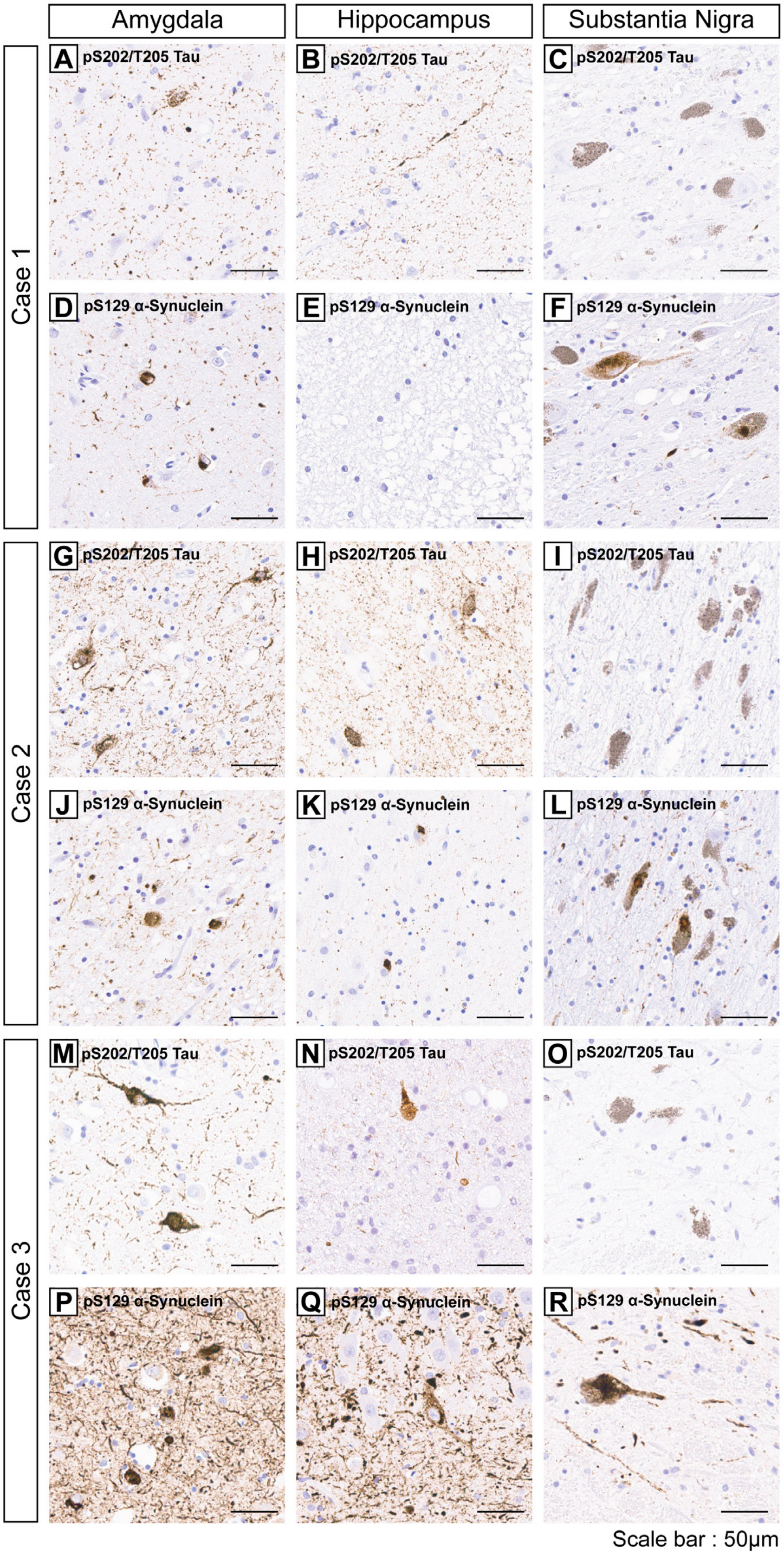

Amygdala
Hippocampus
Substantia Nigra
Case 1
Case 2
Case 3
A pS202/T205 Tau
B pS202/T205 Tau
C pS202/T205 Tau
D pS129 α-Synuclein
E pS129 α-Synuclein
F pS129 α-Synuclein
G pS202/T205 Tau
H pS202/T205 Tau
I pS202/T205 Tau
J pS129 α-Synuclein
K pS129 α-Synuclein
L pS129 α-Synuclein
M pS202/T205 Tau
N pS202/T205 Tau
O pS202/T205 Tau
P pS129 α-Synuclein
Q pS129 α-Synuclein
R pS129 α-Synuclein
Scale bar : 50μm

**Figure S12. Immunohistochemical staining of pS129-α-synuclein and pS202/T205-tau in the hippocampus, amygdala, and substantia nigra of human cases with Lewy body pathology. A-R)** Representative sections from human cases. Amygdala, hippocampus, and substantia nigra sections of case #1 demonstrating staining for pS202/T205-tau **(A-C)** and pS129-α-synuclein **(D-F)**. Sections of case #2 demonstrating staining for pS202/T205-tau **(G-I)** and pS129-α-synuclein **(J-L)** in each brain region. Sections of case #3 demonstrating staining for pS202/T205-tau **(M-O)** and pS129-α-synuclein **(P-R)** in each region. Scale bars: 50 μm.

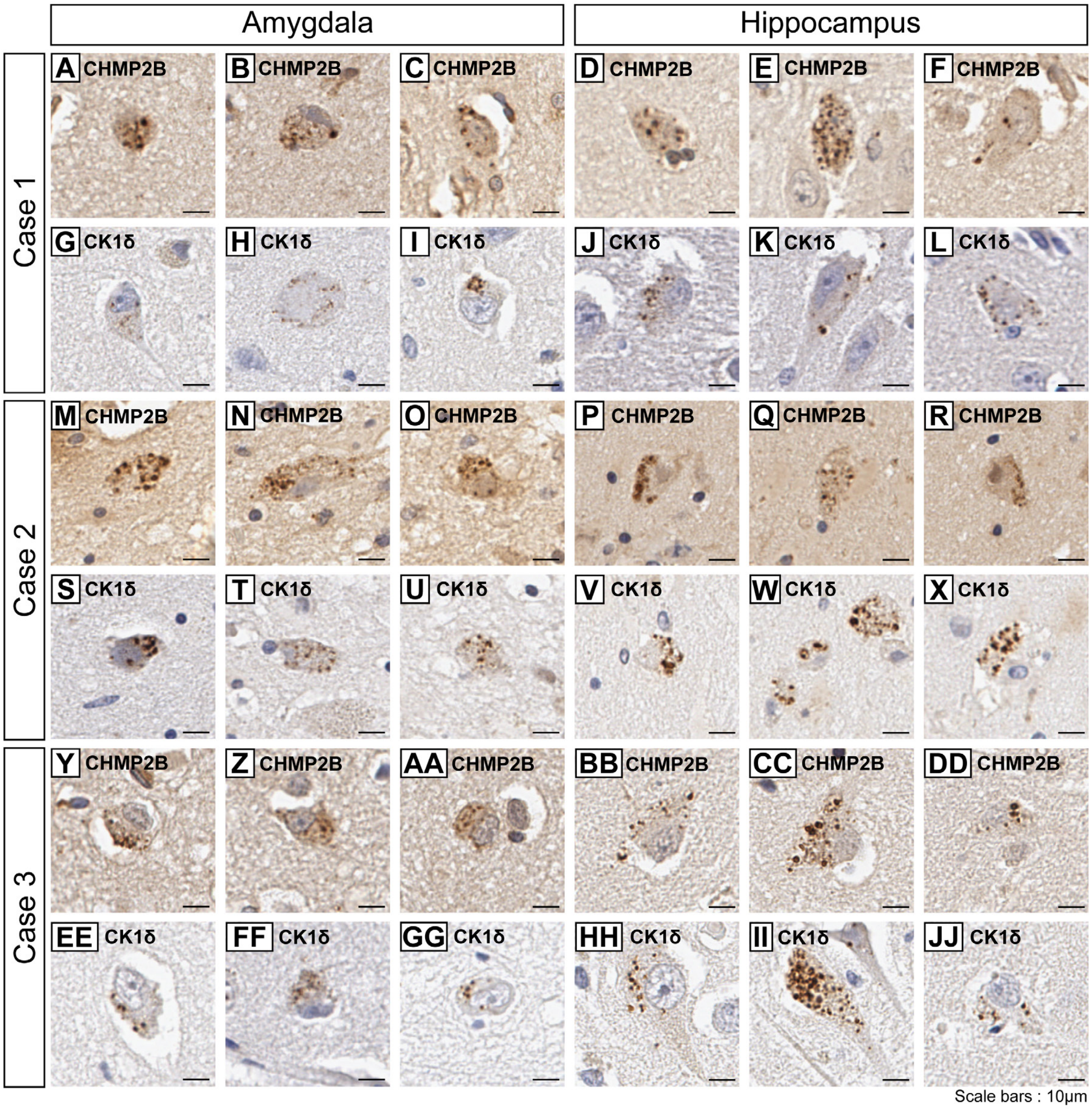


**Figure S13. Immunohistochemical staining of GVB markers in the amygdala and hippocampus of human cases with Lewy body pathology. A-JJ)** Representative sections from human cases depicting IHC staining for GVB markers. Amygdala and hippocampal sections of case #1 demonstrating staining for CHMP2B **(A-F)** and CK1δ **(G-L)**. Sections of case #2 demonstrating IHC for CHMP2B **(M-R)** and CK1δ **(S-X)**. Sections of case #3 demonstrating IHC for CHMP2B **(Y-DD)** and CK1δ **(EE-JJ)**. Scale bars: 10 μm.

| Fluorescent Immunohistochemistry Acquisition Details | | | |
|---|---|---|---|
| Figure | Magnification | Z-step size (μm) | Number of Z steps |
| 4 | 60X | 0.2 | 20 |
| 5 | 60X | 0.5 | 15 |
| 7 | 60X | 0.2 | 14 |
| 8 | 60X | 0.5 | 15 |
| 9 | 60X | 0.2 | 7 |
| S7 | 60X | 0.2 | 20 |
| S8 | 60X | 0.2 | 20 |
| S9 | 60X | 0.2 | 14 |
| S10 | 60X | 0.5 | 15 |
| S11 | 20X | 0.5 | 15 |

**Table S1. Fluorescent Immunohistochemistry Acquisition Details**
